# Mild impairment of mitochondrial function increases longevity and pathogen resistance through ATFS-1-driven activation of p38-regulated innate immunity

**DOI:** 10.1101/2021.03.31.437812

**Authors:** Juliane C. Campos, Ziyun Wu, Paige D. Rudich, Sonja K. Soo, Meeta Mistry, Julio C.B. Ferreira, T. Keith Blackwell, Jeremy M. Van Raamsdonk

## Abstract

While mitochondrial function is essential for life in all multicellular organisms, a mild impairment of mitochondrial function can extend longevity. By understanding the molecular mechanisms involved, these pathways might be targeted to promote healthy aging. In studying two long-lived mitochondrial mutants in *C. elegans,* we found that disrupting subunits of the mitochondrial electron transport chain resulted in upregulation of genes involved in innate immunity, which we found to be dependent on not only the canonical p38-mediated innate immune signaling pathway but also on the mitochondrial unfolded protein response. Both of these pathways are absolutely required for the increased resistance to bacterial pathogens and extended longevity of the long-lived mitochondrial mutants, as is the FOXO transcription factor DAF-16. This work demonstrates that both the p38-mediated innate immune signaling pathway and the mitochondrial unfolded protein response can act on the same innate immunity genes to promote resistance to bacterial pathogens, and that input from the mitochondria can extend longevity by signaling through these two pathways. Combined, this indicates that multiple evolutionarily conserved genetic pathways controlling innate immunity also function to modulate lifespan.

**Significance Statement:** In this work, we explore the relationship between mitochondrial function, aging and innate immunity. We find that mild impairment of mitochondrial function results in upregulation of genes involved in innate immunity, increased resistance to bacterial pathogens and lifespan extension, all of which are dependent on two evolutionarily conserved signaling pathways. This work demonstrates how changes in functional status of the mitochondria can trigger activation of innate immunity, and that the underlying mechanisms are important for the longevity of the organism. This work advances our understanding of connections between metabolism and immunity. As the pathways studied here are conserved up to mammals, these insights may help us to understand the role of mitochondrial health, innate immunity and lifespan in humans.

## Introduction

While aging was long believed to be a stochastic process of damage accumulation, research during the past three decades has demonstrated that lifespan can be strongly influenced by genetics. Single-gene mutations have been shown to extend longevity in model organisms, including yeast, worms, flies and mice. Importantly, genes and interventions that increase lifespan tend to be conserved across species. For example, decreasing insulin-IGF1 signaling, which was first shown to increase lifespan in the worm *C. elegans*(1–3), has subsequently been shown to extend longevity in flies(4) and mice(5), and to be associated with longevity in humans(6). This suggests that studying the aging process in model organisms can provide insights that are relevant to human aging.

The first single gene mutation that was shown to extend lifespan was identified in *C. elegans*(2, 3), and since then this organism has been extensively used to find additional genetic pathways associated with lifespan extension and to elucidate the underlying mechanisms. Among the earliest genes that were shown to influence longevity were genes involved in mitochondrial function. Mutations in *clk-1, nuo-6* and *isp-1* affect different components of the mitochondrial electron transport chain, and all lead to increased lifespan(7–10). In the case of *nuo-6* and *isp-1,* which encode subunits of complex I and complex III respectively, a complete loss of function mutation would be lethal, while point mutations that result in a mild impairment of mitochondrial function extend longevity. Similarly, decreasing the expression of genes involved in mitochondrial function with RNA interference also increases lifespan(11, 12). Importantly, mutations that affect mitochondrial function have also been shown to increase lifespan in other species, including flies(13) and mice(14, 15).

While initially it was believed that the mechanism by which mild impairment of mitochondrial function increased lifespan was through a decrease in the production of reactive oxygen species (ROS) and the resulting oxidative damage(9), more recent studies show that mutations affecting mitochondrial function actually increase the levels of ROS(16). The increase in ROS is required for the long lifespan of these mutants, as treatment with antioxidants can decrease their lifespan to wild-type(16, 17). While multiple factors contributing to the long lifespan of these mitochondrial mutants have been identified(18–23), the precise mechanisms of lifespan extension remain incompletely understood. In our previous work, we have shown that two stress-responsive transcription factors, DAF-16/FOXO3 and ATFS-1/ATF5, are required for the long-lifespan of *nuo-6* and *isp-1* worms(19, 20).

DAF-16/FOXO3 is a FOXO transcription factor that is directly regulated by phosphorylation in response to insulin-IGF1 signaling, a growth factor signaling pathway that begins with the insulin-IGF1 receptor DAF-2. While DAF-16 normally resides in the cytoplasm, when signaling through the insulin-IGF1 pathway is reduced DAF-16 accumulates in nuclei. DAF-16 also translocates to the nucleus in response to various stresses. In the nucleus, DAF-16 upregulates genes involved in stress response and metabolism(24, 25).

Activating transcription factor associated with stress 1 (ATFS-1/ATF5) is the transcription factor that mediates the mitochondrial unfolded protein response (mitoUPR)(26), a stress response pathway that responds to mitochondrial stress in order to restore mitochondrial function. ATFS-1 contains a nuclear localization signal (NLS) and a mitochondrial targeting sequence (MTS). Under normal conditions, ATFS-1 is targeted to the mitochondria where it is imported and degraded. Under conditions of mitochondrial stress, mitochondrial import of ATFS-1 is prevented, resulting in accumulation of ATFS-1 in the cytoplasm, where the NLS translocates ATFS-1 into the nucleus in order to restore mitochondrial homeostasis through alterations in expression of genes involved in protein folding and metabolism(27).

While DAF-16 and ATFS-1 can both contribute to defense against bacterial pathogens(28, 29), the primary innate immune signaling pathway that responds to bacterial pathogen stress is a mitogen-activated protein kinase (MAPK) signaling pathway, which has been found to be conserved from invertebrates to mammals(30–32). In this pathway, NSY-1/ASK1 (MAPK kinase kinase) signals to SEK-1/MKK3/MKK6 (MAPK kinase), which signals to PMK-1/p38 (MAPK) (33, 34) (**Fig. S1**). Downstream of this pathway, the transcription factor ATF-7/ATF2/ATF7/CREB5 acts to modulate the expression of genes involved in innate immunity (35, 36). While ATF-7 normally acts as a repressor of gene function, when it is phosphorylated by PMK-1, ATF-7 functions as an activator of p38/ATF-7-regulated immunity gene expression (**Fig. S1**).

In this work, we show that the p38-mediated innate immune signaling pathway is essential for the longevity and pathogen resistance of the long-lived mitochondrial mutants, *nuo-6* and *isp-1.* We find that both strains exhibit an upregulation of genes involved in innate immunity that is driven by the activation of the mitoUPR, but also dependent on the p38-mediated innate immune signaling pathway, leading to an increased resistance to bacterial pathogens. The p38-mediated innate immune signaling pathway is absolutely required for the long lifespan of *nuo-6* and *isp-1* mutants. Finally, we demonstrate that activation of the mitoUPR is sufficient to upregulate innate immunity genes, and is also required for their upregulation in *nuo-6* mutants. Overall, this work demonstrates the importance of the mitoUPR in upregulating innate immunity in response to signals from the mitochondria, and delineates a clear role of innate immune signaling pathways in determining lifespan.

## Results

### Long-lived mitochondrial mutants exhibit broad upregulation of genes involved in innate immunity that is dependent on p38-mediated innate immune signaling pathway

While mild impairment of mitochondrial function has been shown to extend longevity, the underlying mechanisms are yet to be fully elucidated. When mitochondrial function is impaired, mitochondria are able to communicate with the nucleus to alter nuclear gene expression. To obtain a comprehensive, unbiased view of the transcriptional changes that result from impairment of mitochondrial function, we used RNA sequencing (RNA-seq) to examine gene expression in two long-lived mitochondrial mutants, *nuo-6* and *isp-1*. After determining which genes were differentially expressed compared to wild-type worms, we identified groups of genes that showed enrichment. Among the genes that showed enrichment were genes involved in innate immunity. These genes encode proteins that function to inhibit the growth and survival of pathogenic bacteria, and to repair or remove damage to the worm(37, 38). Accordingly, we decided to investigate the role of innate immunity in the long lifespan of these long-lived mitochondrial mutants.

To determine the extent to which genes involved in innate immunity are upregulated in the long-lived mitochondrial mutants, *nuo-6* and *isp-1,* we first examined eight genes, which others have used to monitor innate immune activity(29, 35, 37, 39–41). These genes included *T24B8.5, K08D8.5, F55G11.8, clec-65, clec-67, dod-22, Y9C9A.8* and *C32H11.4*. All of these genes are upregulated in response to exposure to the bacterial pathogen *Pseudomonas aeruginosa* strain PA14 and five of eight have been shown to be direct targets of the p38-mediated innate immune signaling pathway in a ChIP-seq analysis of ATF-7(36). In examining the expression of these genes in our RNA-seq data, we found that all eight genes were significantly upregulated in both *nuo-6* and *isp-1* worms (**Fig. 1A**).

**Figure 1.**
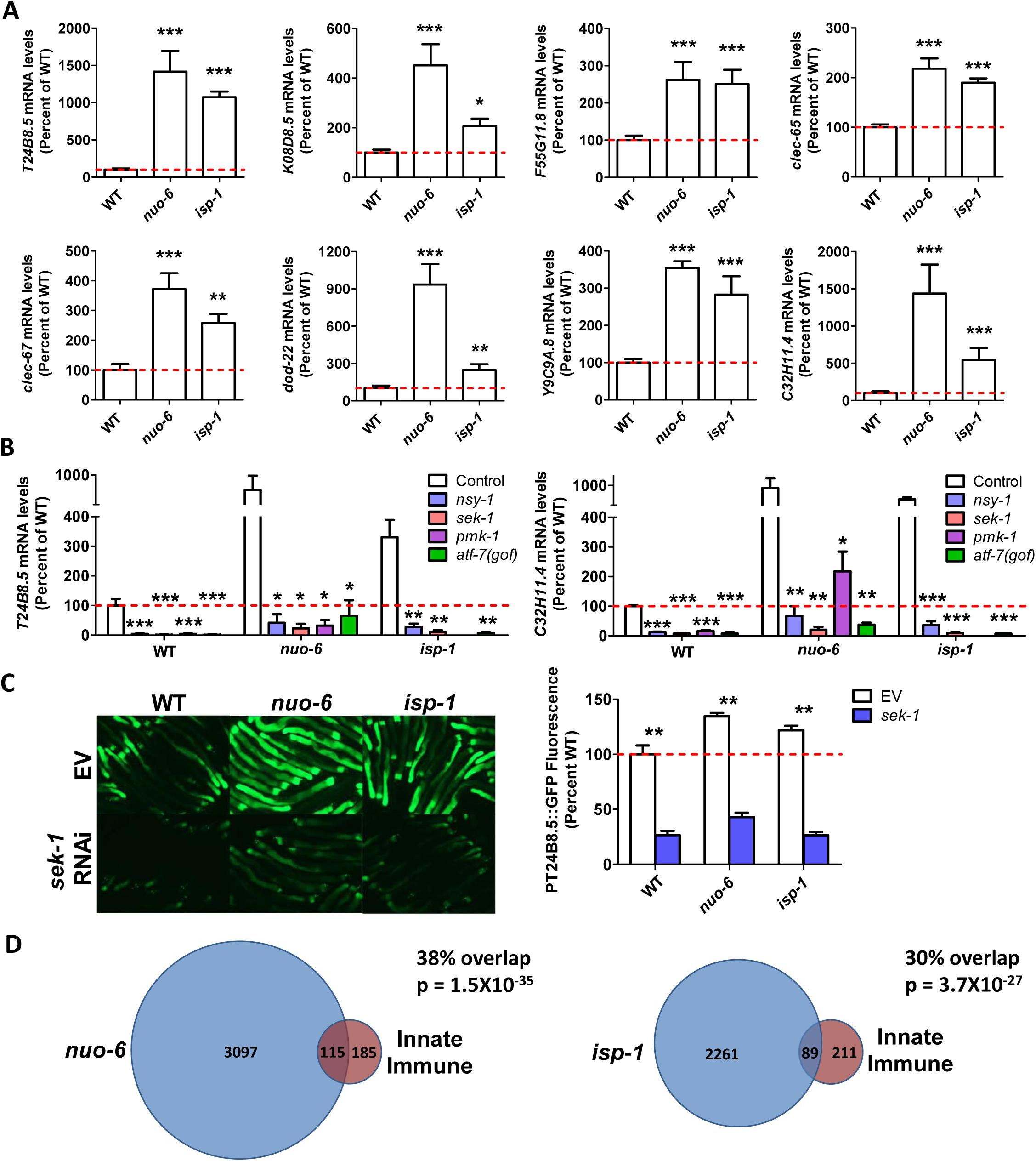
Innate immunity genes are upregulated in long-lived mitochondrial mutants. **A.** Genes involved in innate immunity are significantly upregulated in *nuo-6* and *isp-1* worms. Gene expression changes in the mitochondrial mutants were determined by RNA sequencing of six biological replicates and compared to wild-type N2 worms. Results represent counts per million (CPM) expressed as a percentage of wild-type. **B.** Upregulation of innate immunity genes is dependent on the p38-mediated innate immune signaling pathway including NSY-1, SEK-1, PMK-1 and ATF-7. Gene expression changes were examined by quantitative RT-PCR with 3 biological replicates per strain. **C.** Using a fluorescent reporter strain for the innate immunity gene *T24B8.5* confirms that innate immunity genes are upregulated in the long-lived mitochondrial mutants and that components of the p38-mediated innate immune signaling pathway are required for their upregulation. **D.** Differentially expressed genes in long-lived mitochondrial mutants show significant overlap with genetic targets of the p38-mediated innate immune signaling pathway. Significantly modulated genes in long-lived mitochondrial mutants *nuo-6* and *isp-1* (six biological replicate per strain) were compared to genes that are modulated in response to exposure to the bacterial pathogen *P. aeruginosa* strain PA14 in a PMK-1- and ATF-7-dependent manner as identified by Fletcher et al., 2019. There is a highly significantly degree of overlap between genes upregulated by activation of the p38-mediated innate immune pathway and genes upregulated in *nuo-6* and *isp-1* mutants. Similarly, there is a highly significantly degree of overlap between genes downregulated by activation of the p38-mediated innate immune pathway and genes downregulated in *nuo-6* and *isp-1* mutants. Error bars indicate SEM. *p<0.05, **p<0.01, ***p<0.001.

To determine whether the upregulation of innate immunity genes in the long-lived mitochondrial mutants requires the p38-mediated innate immune signaling pathway (NSY-1 → SEK-1 → PMK-1 → ATF-7), we generated double mutants of *nuo-6* and *isp-1* with all of these genes before measuring gene expression with quantitative RT-PCR (qPCR). We used loss of function mutants for *nsy-1, sek-1* and *pmk-1.* Since ATF-7 normally acts as a repressor, we used the *qd22* gain-of-function mutation for this gene. This mutation prevents phosphorylation of ATF-7 by PMK-1 thereby making the mutant ATF-7 a constitutive repressor(35) (**Fig. S1**). As in the RNA-seq data, the results from the qPCR experiments showed that innate immune genes are upregulated in *nuo-6* and *isp-1* worms. Importantly, we found that in both *nuo-6* and *isp-1* worms that disruption of genes involved in the p38-mediated innate immune signaling pathway (*nsy-1, sek-1, pmk-1, atf-7(gof)*) prevented the upregulation of innate immunity genes (**Fig. 1B, Fig. S2**).

To confirm these results using an alternative approach, we crossed a fluorescent reporter strain for one of the innate immunity genes (*T24B8.5*)(35) to *nuo-6* and *isp-1* mutants and examined the effect of knocking down *sek-1* through RNA interference (RNAi). Again, using this approach, we found that *T24B8.5* is upregulated in *nuo-6* and *isp-1* worms, and that this upregulation is dependent on SEK-1 (**Fig. 1C**). Combined, these results demonstrate that innate immunity genes are upregulated in the long-lived mitochondrial mutants, *nuo-6* and *isp-1,* and that this upregulation is dependent on the p38-mediated innate immune signaling pathway.

To further examine the expression of innate immunity genes in *nuo-6* and *isp-1* mutants, we compared the differentially expressed genes in these mutants to a more comprehensive and unbiased list of genes involved in innate immunity. A recent study defined the changes in gene expression that result from exposure to the bacterial pathogen *Pseudomonas aeruginosa* strain PA14 and determined which of these changes in gene expression are dependent on PMK-1 and ATF-7(36). In total, they reported 300 genes that were upregulated by exposure to PA14 in a PMK-1 and ATF-7-dependent manner, and 230 genes that were downregulated by exposure to PA14 in a PMK-1 and ATF-7-dependent manner.

We compared these lists of PA14-modulated, PMK-1-dependent, ATF-7-dependent genes to genes that we found to be significantly upregulated or downregulated in *nuo-6* and *isp-1* mutants. We found that of the genes that are upregulated by PA14 exposure in a PMK-1 and ATF-7-dependent manner, 38% (p=1.5 x 10^-35^) and 30% (p=3.7 x 10^-27^) are also upregulated in *nuo-6* and *isp-1* mutants, respectively (**Fig. 1D**). There is a high degree of overlap between genes upregulated in *nuo-6* and *isp-1* mutants (71%), and this includes the genes involved in innate immunity (**Table S1**; 71 overlapping genes of 89/115 innate immunity genes upregulated in *nuo-6* or *isp-1,* respectively). In contrast, there was no significant overlap of the same genes upregulated by PA14 exposure with genes downregulated in *nuo-6* and *isp-1* mutants. In examining the list of genes that are downregulated by exposure to PA14 in a PMK-1 and ATF-7-dependent manner, we found that 20% (p=8.4 x 10^-7^) and 20% (p=7.5 x 10^-5^) were also downregulated in *nuo-6* and *isp-1* mutants, respectively.

To more comprehensively compare the PA14-modulated, PMK-1-dependent, ATF-7-dependent gene expression changes to gene expression changes in *nuo-6* and *isp-1* mutants, we generated heat maps comparing the expression of PA14-modulated, PMK-1-dependent, ATF-7-dependent genes between wild-type worms and *nuo-6* or *isp-1* mutants. Among the genes that are upregulated by PA14 exposure in a PMK-1 and ATF-7-dependent manner, many of these genes are upregulated in *nuo-6* and *isp-1* mutants compared to wild-type worms, while a small subset show decreased expression (**Figs. S3,S4**). Among the genes that are downregulated by PA14 exposure in a PMK-1 and ATF-7-dependent manner, many of these genes are downregulated in *nuo-6* and *isp-1* mutants compared to wild-type worms, while a number of these genes also show increased expression (**Figs. S5,S6**).

Overall, these results indicate that a mild impairment of mitochondrial function caused by mutations in *nuo-6* or *isp-1* leads to a broad upregulation of genes involved in innate immunity that is dependent on the p38-mediated innate immune signaling pathway. At the same time, a smaller proportion of these genes show decreased expression suggesting that the correct balance of innate immune gene expression may be required to optimize stress resistance and lifespan.

### Long-lived mitochondrial mutants have increased resistance to bacterial pathogens, which requires the p38-mediated innate immune signaling pathway

Based on our observation that the long-lived mitochondrial mutants, *nuo-6* and *isp-1,* have increased expression of many genes involved in innate immunity, we next sought to determine whether this increase in their expression resulted in enhanced resistance to bacterial pathogens. To test pathogen resistance, we exposed worms to PA14 in a slow kill assay where worms die from the ingestion and internal proliferation of the PA14 bacteria(42–44). We found that both *nuo-6* and *isp-1* worms exhibited significantly increased survival on PA14 bacteria compared to wild-type worms (**Fig. 2A**). The increase in survival of *nuo-6* and *isp-1* worms did not result from reduced exposure to the pathogenic bacteria as their tendency to avoid PA14 was equivalent to wild-type worms (**Fig. S7**).

**Figure 2.**
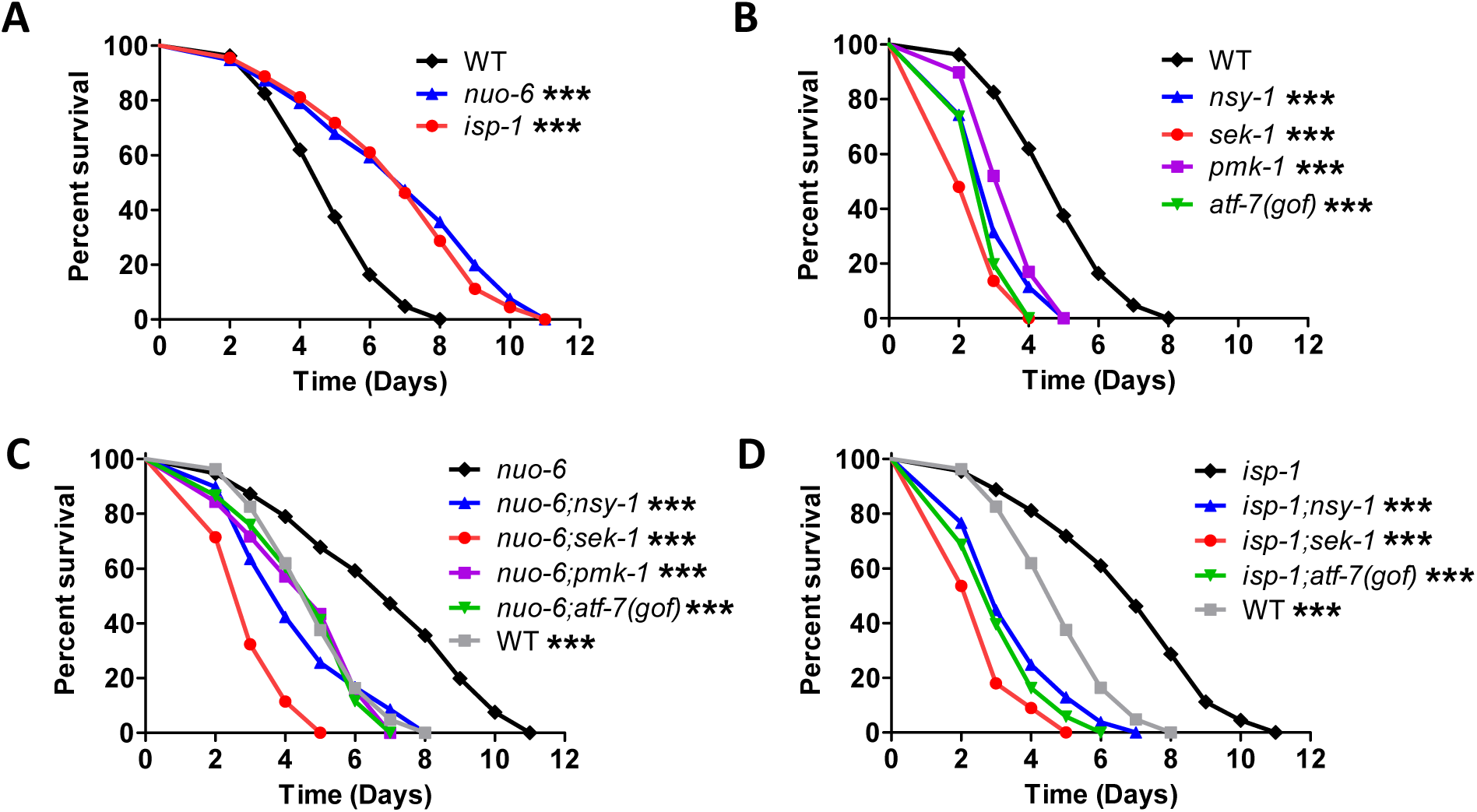
Long-lived mitochondrial mutants exhibit increased resistance to bacterial pathogens that is dependent on presence of p38-mediated innate immune signaling pathway. Resistance to bacterial pathogens was tested by exposing worms to *Pseudomonas aeruginosa* strain PA14 in a slow kill assay. **A.** Both *nuo-6* and *isp-1* long-lived mitochondrial mutants showed increased resistance compared to wild-type worms. Disruption of components of the p38-mediated innate immune signaling pathway (*nsy-1, sek-1, pmk-1, atf-7*) markedly decreases resistance to PA14 in wild-type (**B**), *nuo-6* (**C**) and *isp-1* (**D**) worms. Statistical analysis on survival plots was performed with log-rank test. Significance is indicated between the strain listed on top and all other strains. ***p<0.001. All strains were tested in a single parallel experiment. Data from panel A is repeated in panels B, C, and D for direct comparison. Raw data and N for each strain can be found in **Table S2**.

To determine the extent to which their enhanced resistance to bacterial pathogens is dependent on the p38-mediated innate immune signaling pathway, we next examined PA14 resistance in *nuo-6* and *isp-1* worms in which genes in this pathway were disrupted. In wild-type worms, mutations in *nsy-1, sek-1, pmk-1* or *atf-7(gof)* significantly decrease the survival of worms exposed to PA14 (**Fig. 2B**). Similarly, we found that disruptions of these innate immune signaling genes in *nuo-6* (**Fig. 2C**) or *isp-1* (**Fig. 2D**) worms also results in a significant decrease in survival on PA14 bacteria. In *nuo-6* worms, survival was decreased back to wild-type by mutations in *nsy-1, pmk-1* or *atf-7(gof)*, while a larger decrease in survival was observed with the *sek-1* mutation. In *isp-1* worms, mutations in *nsy-1, sek-1* and *atf-7(gof)* all decreased survival to a greater extent than in *nuo-6* mutants, and in each case the survival of the double mutant was less than in wild-type worms (**Fig. S8**). Combined, this shows that *nuo-6* and *isp-1* worms have increased resistance to bacterial pathogens, which is dependent on the p38-mediated innate immune signaling pathway.

### p38-mediated innate immune signaling pathway is required for extended longevity in long-lived mitochondrial mutants

The p38-mediated innate immune signaling pathway is required for the lifespan extension resulting from decreased insulin-IGF1 signaling (*daf-2* mutants) and dietary restriction(38, 39). Interestingly, in both *daf-2* mutants and dietary restricted worms expression of p38-regulated innate immunity genes is largely decreased, and activation of this pathway to higher levels is deleterious for lifespan extension, suggesting that a lower level of immune activation is optimal under these conditions(39). We decided to examine the role of the p38-mediated innate immune signaling pathway in the long lifespan of *nuo-6* and *isp-1* mutants, in which we found these immunity genes were largely upregulated. To do this, we genetically disrupted components of the p38-mediated innate immune signaling pathway in *nuo-6* and *isp-1* mutants and quantified the resulting effect on lifespan. In *nuo-6* mutants, we found that mutation of *nsy-1, sek-1, pmk-1* or *atf-7(gof)* reduced the long lifespan of *nuo-6* worms to near wild-type lifespan, almost completely preventing any increase in lifespan resulting from the *nuo-6* mutation (**Fig. 3A-D**). Similarly, we found that mutations in the p38-mediated innate immune signaling pathway also markedly reduced the lifespan of *isp-1* worms (**Fig. 3E-H**). Mutations in *nsy-1, sek-1,* or *atf-7(gof)* decreased *isp-1* lifespan to near wild-type lifespan, while *pmk-1* RNAi reduced the lifespan extension in *isp-1* worms by half (we were unable to generate *isp-1 pmk-1* double mutants because of the proximity of the two genes on the same chromosome).

**Figure 3.**
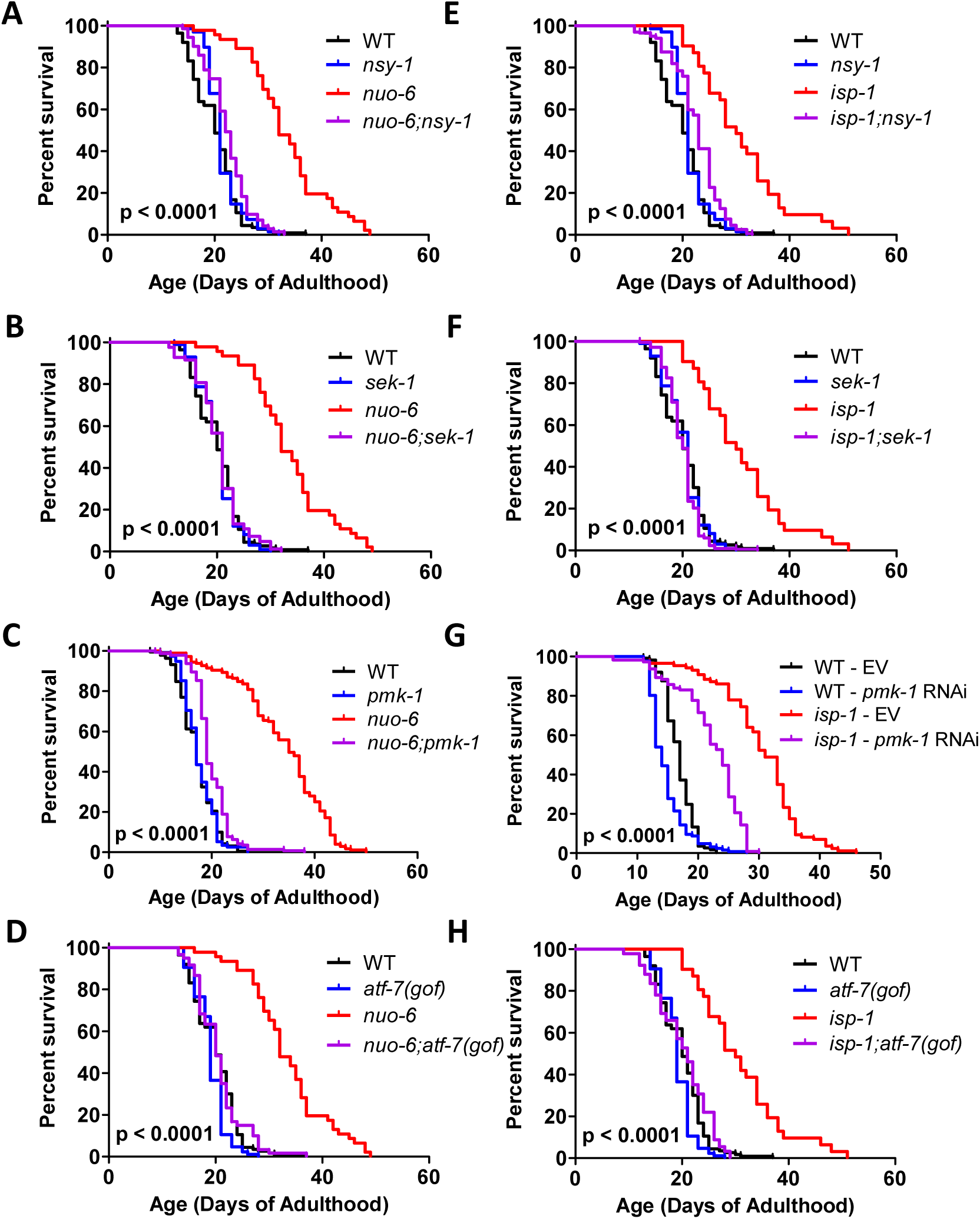
Disruption of the p38-mediated innate immune signaling pathway markedly ablates extended longevity of long-lived mitochondrial mutants. To examine the role of the p38-mediated innate immune signaling pathway in the long-lifespan of the *nuo-6* and *isp-1* mitochondrial mutants, we crossed the long-lived mitochondrial mutants to worms with mutations in *nsy-1, sek-1, pmk-1* and *atf-7(gof).* In each case, mutation of genes involved in p38-mediated innate immunity signaling markedly reduced the lifespan of the long-lived mitochondrial mutant but had little or no impact on wild-type lifespan. Results are from a minimum of three biological replicates with at least 40 worms per replicate. Statistical analysis on survival plots was performed with log-rank test. p-value indicate significance between red and purple lines. The lifespan results for panels A, B, D, E, F, and H were performed in a single parallel experiment. Control strains are shown in multiple panels for direct comparison. Raw data and N for each strain can be found in **Table S2**.

Although OP50 bacteria, which is typically used for *C. elegans* maintenance and lifespan assays, is considered to be non-pathogenic, it can have detrimental effects on lifespan through proliferation in the pharynx and intestine (45, 46). Accordingly, we also measured the lifespan of *nuo-6* and *isp-1* mutants on non-proliferating OP50 bacteria, as well as the dependency of their long lifespan on the p38-mediated innate immunity pathway under these conditions. Similar to what we observed on live bacteria, we found that *nuo-6* and *isp-1* mutants are long-lived on non-proliferating bacteria, and that their long lifespan is still completely dependent on the p38-mediated innate immune signaling pathway (**Fig S9**).

Combined, these results clearly indicate that the p38-mediated innate immune signaling pathway is required for the extended longevity in the long-lived mitochondrial mutants, *nuo-6* and *isp-1.* However, the fact that effect of disrupting the p38-innate immune signaling pathway on *nuo-6* and *isp-1* lifespan is independent of bacterial proliferation suggests that the lifespan extension resulting from mild impairment of mitochondrial function is not necessarily a direct consequence of their enhanced resistance to bacterial pathogens, but more likely a result of other functions of the p38-mediated innate immune signaling pathway.

### p38-mediated innate immune signaling pathway is not activated in long-lived mitochondrial mutants

Since p38 signaling can be activated by ROS(47, 48), and ROS levels are increased in *nuo-6* and *isp-1* mutants(16), we hypothesized that the upregulation of innate immunity genes in these mutants resulted from increased signaling through the p38-mediated innate immune signaling pathway. To test this, we examined the extent to which this pathway might be activated in these mutants by quantifying the ratio of phosphorylated (active) p38/PMK-1 to total p38/PMK-1 by Western blotting(47). To validate this approach, we showed that decreasing innate immune signaling with *sek-1* RNAi and increasing innate immune signaling with *vhp-1* RNAi (VHP-1 is a phosphatase that inhibits PMK-1/p38 signaling) alters the ratio of phospho-p38 to p38 as expected (**Fig. S10**).

Having shown that known modulators of PMK-1/p38 activation alter PMK-1/p38 phosphorylation in a predictable manner, we examined PMK-1/p38 activation in *nuo-6* and *isp-1* mutants. In both cases we found that neither the levels of PMK-1/p38 protein nor the proportion of PMK-1/p38 protein that is phosphorylated are increased compared to wild-type (**Fig. 4A,B**). Thus, even though *nuo-6* and *isp-1* mutants have increased expression of ATF-7 target genes, there is no detectable increase in PMK-1/p38 activation. Since the p38-mediated innate immune signaling pathway is required for the increased expression of ATF-7 target genes even though p38 signaling is not increased, this suggests that this pathway is playing a permissive role, and that other mechanisms are driving the innate immune activation in the long-lived mitochondrial mutants.

**Figure 4.**
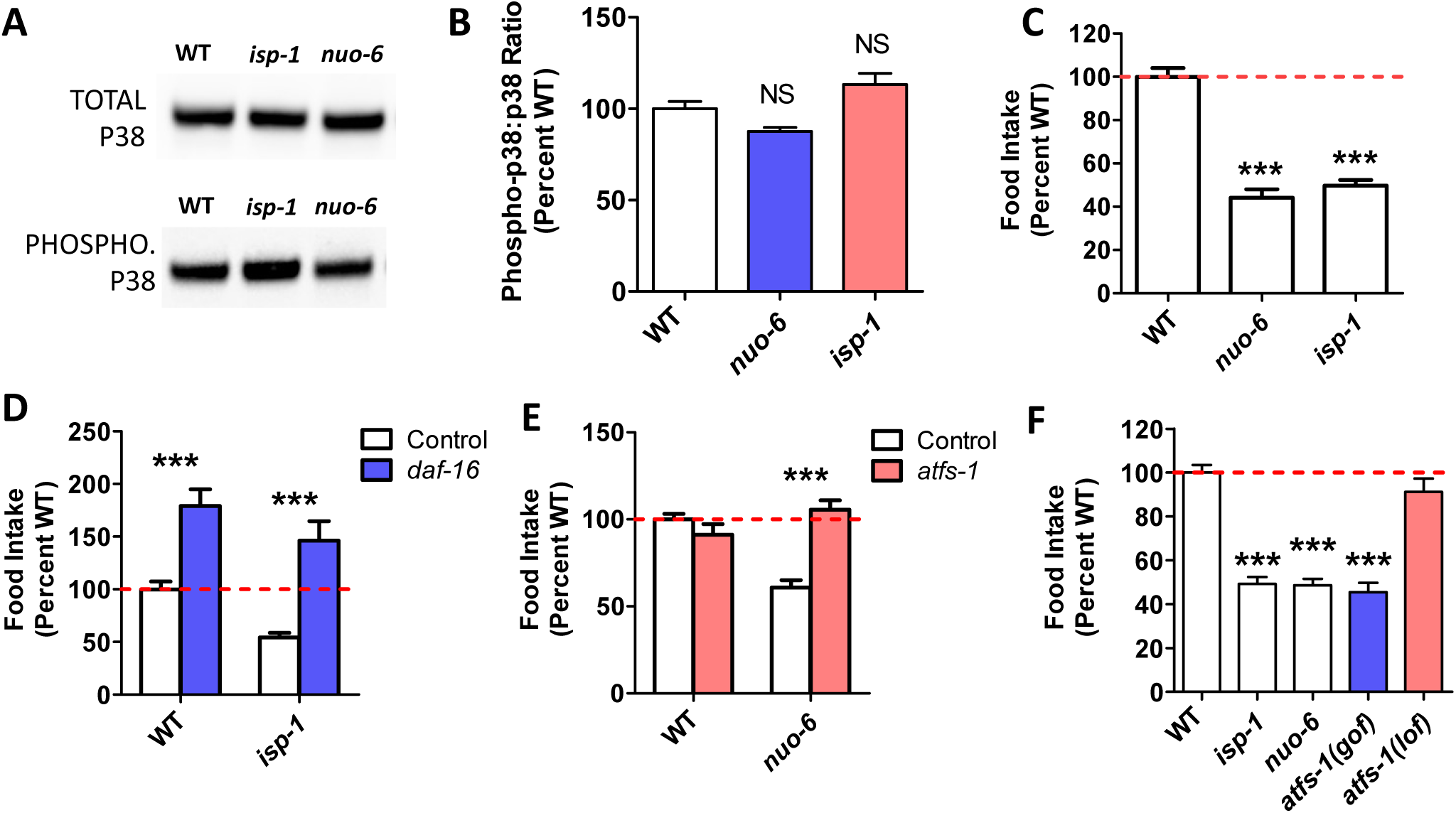
p38-mediated innate immune signaling pathway is not activated in long-lived mitochondrial mutants. **A.** To determine the extent to which the p38-mediated innate immunity pathway is activated in *nuo-6* and *isp-1* mutants, we measured the ratio of phosphorylated p38/PMK-1 to total p38/PMK-1 by Western blotting. **B.** There was no difference in the ratio of phosphorylated p38/PMK-1 to total p38/PMK-1 in the long-lived mitochondrial mutants compared to wild-type. Four biological replicates per strain were quantified. See **Supplemental Data** for complete images of Western blots including loading controls. **C.** Both long-lived mitochondrial mutants, *nuo-6* and *isp-1,* exhibit markedly decreased food consumption compared to wild-type worms. At least six biological replicates per strain were measured, each including at least 3 technical replicates. **D.** Deletion of *daf-16* significantly increases food consumption in both wild-type and *isp-1* mutants. Three biological replicates per strain were measured. **E.** While disruption of *atfs-1* does not affect food intake in wild-type worms, it increases food consumption in *nuo-6* worms back to wild-type. *atfs-1* loss of function allele was *gk3094.* Three biological replicates per strain were measured. **F.** Constitutive activation of *atfs-1* is sufficient to decrease food consumption to the same level as in long-lived mitochondrial mutants. Three biological replicates per strain were measured. *atfs-1(lof)* allele is *gk3094, atfs-1(gof)* allele is *et17.* Error bars indicate SEM. **p<0.01, ***p<0.001.

Since the p38-mediated innate immune signaling pathway can be activated by nutrient consumption independently of pathogen exposure(39), we wondered whether altered feeding affects the level of activation of this pathway in the long-lived mitochondrial mutants, countering the effects of elevated ROS. Accordingly, we quantified food consumption in *nuo-6* and *isp-1* worms directly by monitoring the decrease in OP50 bacteria from day 3 to day 12 of adulthood(39). We found that both long-lived mutants show a dramatic decrease in food consumption compared to wild-type worms (**Fig. 4C**). This decrease in food consumption does not result from a difference in occupancy of the bacteria as *nuo-6* and *isp-1* worms spend the amount of time on the bacteria as wild-type worms (**Fig. S7**). The decrease in food consumption in *nuo-6* and *isp-1* worms would be predicted to decrease p38-mediated innate immune signaling, and might be sufficient to decrease any ROS-induced activation of this pathway in these mutants. The fact that the activation of the p38-mediated innate immune signaling pathway is not decreased in the long-lived mitochondrial mutants, and that target genes of this pathway are upregulated, again indicates that other factors are acting in parallel to p38 signaling to upregulate the expression of innate immunity genes.

### Decreased feeding in long-lived mitochondrial mutants is mediated by activation of the mitoUPR

We next wanted to define the mechanism by which feeding is decreased in the long-lived mitochondrial mutants. Since DAF-16 is activated in *nuo-6* and *isp-1* mutants (20) and disruption of *daf-16* has been shown to increase food intake in wild-type and *daf-2* worms(39), we sought to determine whether *daf-16* is required for the decreased food intake observed in the long-lived mitochondrial mutants. We found that deletion of *daf-16* increased food consumption in both wild-type and *isp-1* worms (**Fig. 4D**), thereby indicating that DAF-16 is required for the decreased food consumption in *isp-1* mutants.

Since activation of ATFS-1 is sufficient to upregulate DAF-16 target genes(19) and ATFS-1 is activated in the long-lived mitochondrial mutants(19), we wondered whether ATFS-1 also contributes to the decreased food intake in the long-lived mitochondrial mutants. While disruption of *atfs-1* (using the *gk3094* deletion mutation) had no effect on feeding in a wild-type background, loss of *atfs-1* reverted food consumption in *nuo-6* worms to wild-type (**Fig. 4E**). This indicates that both *daf-16* and *atfs-1* are required for the decreased food intake in the long-lived mitochondrial mutants.

Having shown that activation of ATFS-1 contributes to the decreased food intake in *nuo-6* mutants, we wondered whether activation of ATFS-1 alone is sufficient to decrease food consumption. To test this idea, we measured food consumption in a constitutively activated *atfs-1* mutant (*et17*). In this mutant, the mitochondrial targeting sequence of ATFS-1 is disrupted resulting in nuclear localization and constitutive activation of ATFS-1 target genes(49). We found that the constitutively activated *atfs-1* mutant *et17* showed a marked decrease in food consumption, equivalent to that observed in *nuo-6* and *isp-1* mutants (**Fig. 4F**). Combined, this indicates that activation of ATFS-1 is sufficient to decrease food consumption, and is required for the diminished food intake in *nuo-6* worms.

### The FOXO transcription factor DAF-16 is required for bacterial pathogen resistance in long-lived mitochondrial mutants but does not account for activation of innate immunity genes

As we have previously shown that target genes of the FOXO transcription factor DAF-16 are upregulated in the long-lived mitochondrial mutants(20), and others have shown that DAF-16 is required for the enhanced bacterial pathogen resistance of long-lived *daf-2* mutants(38), we next examined the contribution of DAF-16 to bacterial pathogen resistance in long-lived mitochondrial mutants. We found that disruption of *daf-16* completely abolished the increased resistance to bacterial pathogens in *isp-1* worms (**Fig. 5A**), thereby indicating that DAF-16 is required for the enhanced bacterial pathogen resistance in *isp-1* worms.

**Figure 5.**
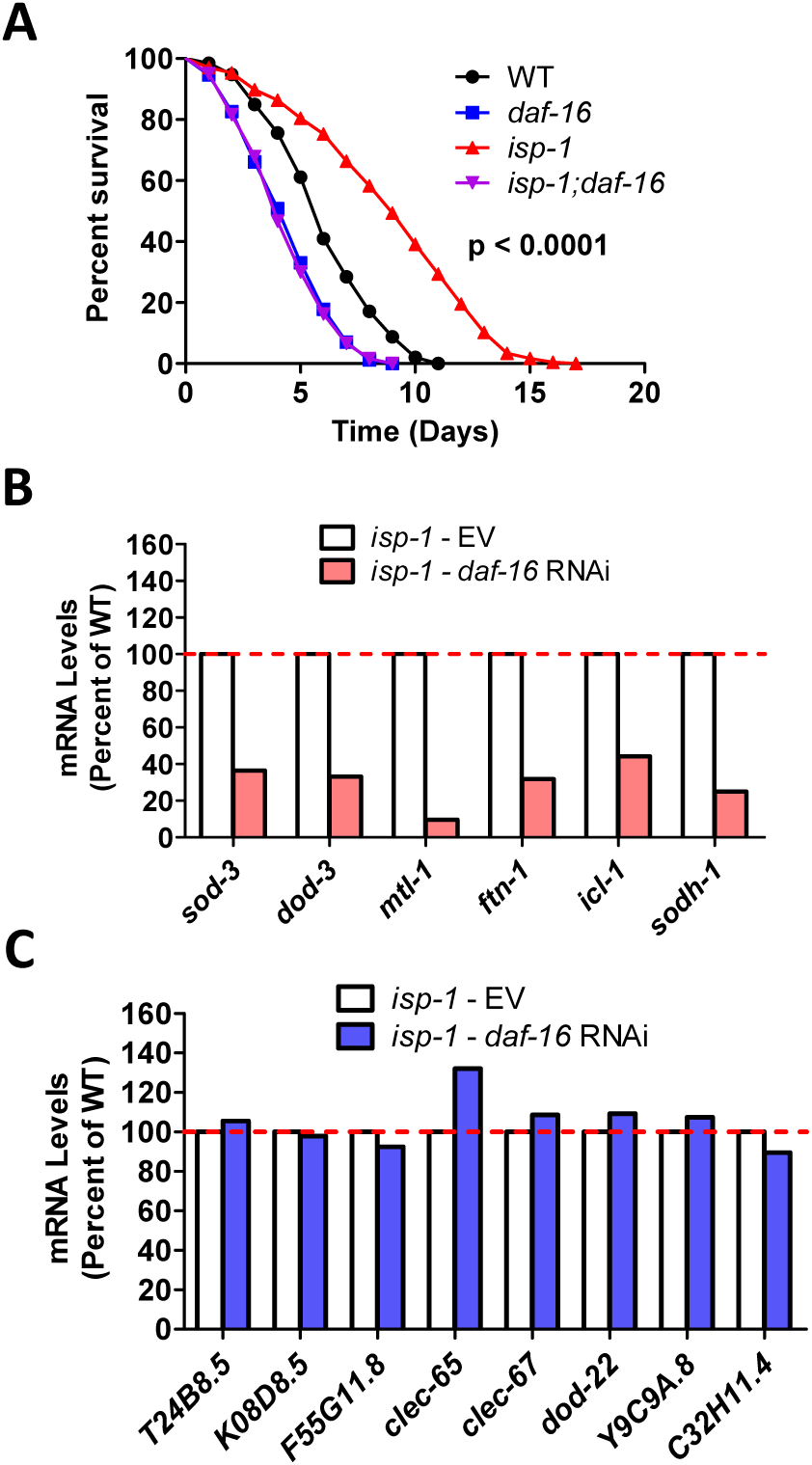
DAF-16/FOXO is required for increased resistance to bacterial pathogens in long-lived mitochondrial mutants. **A.** A deletion of *daf-16* completely abolishes the increased resistance to bacterial pathogens in *isp-1* mutants. p-value indicates the significance of difference between red and purple lines. Three biological replicates per strain were measured. While *daf-16* RNAi effectively decreased the expression of DAF-16 target genes (**B**, *sod-3, dod-3, mtl-1, ftn-1, icl-1, sodh-1*), it did not affect the expression of any of the innate immunity genes (**C**, *T24B8.5, K08D8.5, F55G11.8, clec-65, clec-67, dod-22, Y9C9A.8* and *C32H11.4*). This indicates that DAF-16 is not required for expression of innate immune signaling pathway target genes in *isp-1* mutants. *daf-16* expression was knocked down using RNAi beginning at the L4 stage of the parental generation. RNA was isolated from six biological replicates at the young adult stage of the experimental generation. RNA from the six biological replicates was pooled for RNA sequencing.

To determine the extent to which the effects of DAF-16 on bacterial pathogen resistance might be mediated through the same genes that are modulated by the p38-mediated innate immune signaling pathway, we examined the effect of *daf-16* RNAi on the expression of innate immune genes in wild-type, *nuo-6* and *isp-1* worms. While *daf-16* RNAi was effective in decreasing the expression of DAF-16 target genes (*sod-3, dod-3, mtl-1, ftn-1, icl-1, sodh-1*) in all three strains, there was little or no effect on genes involved in innate immunity (*T24B8.5, K08D8.5, F55G11.8, clec-65, clec-67, dod-22, Y9C9A.8* and *C32H11.4*) (**Fig. 5B,C; Fig. S11**). This finding is consistent with a previous study that compared microarray data of genes modulated by DAF-16 and PMK-1 and observed no overlap, despite both pathways being required for pathogen resistance in *daf-2* mutants(38). Thus, although DAF-16 activation contributes to the enhanced pathogen resistance in the long-lived mitochondrial mutants, it does not contribute to the upregulation of p38/ATF-7-regulated innate immunity genes in these worms.

### The mitoUPR transcription factor ATFS-1 mediates upregulation of innate immune genes in long-lived mitochondrial mutants

In our previous studies, we have shown that the mitoUPR is activated in *nuo-6* and *isp-1* worms, and that this activation is required for their increased lifespan(19). As activation of the mitoUPR by RNAi against *spg-7* (the worm homolog of SPG7/paraplegin) has been shown to increase resistance to PA14 and upregulate innate immunity genes(29), we hypothesized that the ATFS-1 activation that occurs in *nuo-6* and *isp-1* mutants(19) might contribute to their enhanced resistance to PA14.

To test this, we examined the effect of inhibiting the mitoUPR on the survival of *nuo-6* worms under bacterial pathogen stress. We found that disruption of *atfs-1* completely ablates the bacterial pathogen resistance of *nuo-6* worms such that *nuo-6;atfs-1* worms have decreased survival compared to wild-type or *atfs-1* single mutants (**Fig. 6A**). This indicates that activation of the mitoUPR is required for the enhanced pathogen resistance in *nuo-6* mutants.

**Figure. 6.**
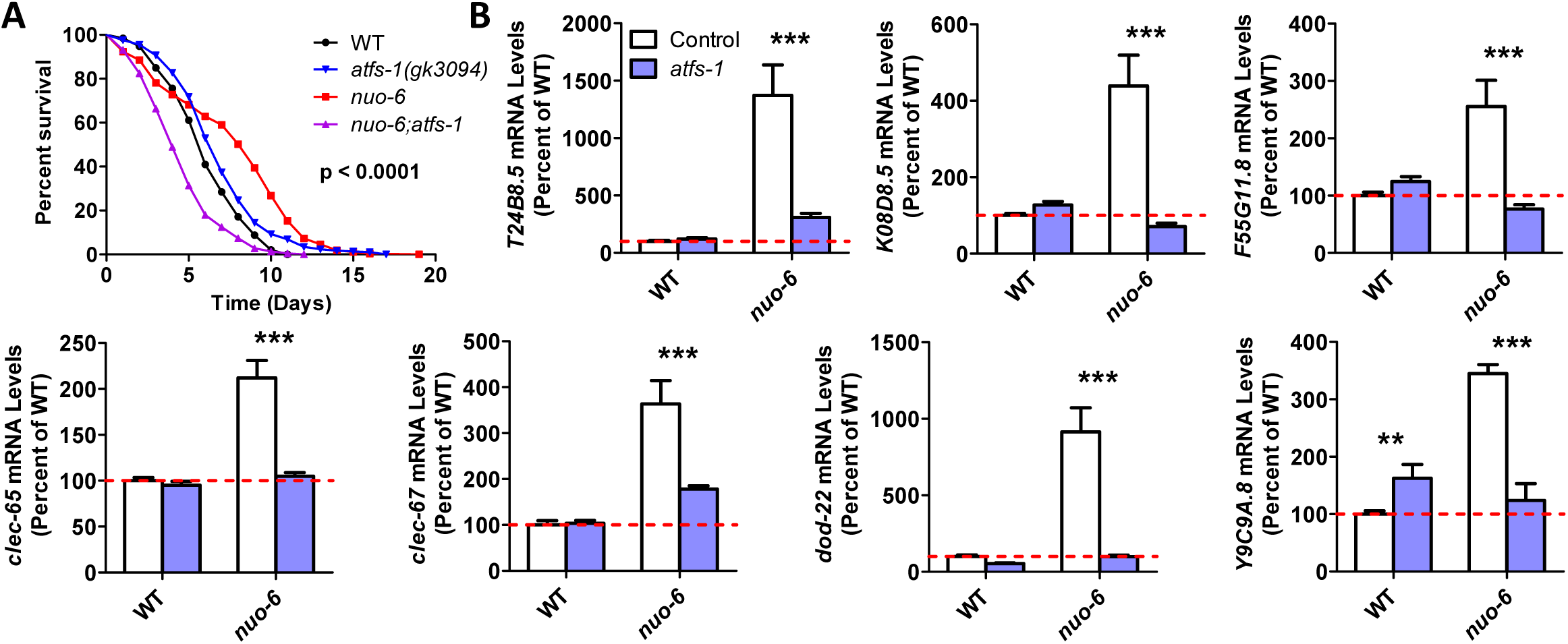
ATFS-1 is required for increased bacterial pathogen resistance and upregulation of innate immunity genes in long-lived *nuo-6* mitochondrial mutants. **A.** To examine the role of the ATFS-1 transcription factor and the mitochondrial unfolded protein response in the enhanced bacterial pathogen resistance in *nuo-6* mutants, we examined the effect of disrupting *atfs-1* in *nuo-6* mutants. Loss of *atfs-1* completely abolished the increased resistance to bacterial pathogens in *nuo-6* worms. p-value indicates significance of difference between red and purple lines. *nuo-6;atfs-1* contains the *gk3094* deletion allele. Three biological replicates per strain were measured. **B.** To examine the role of ATFS-1 in the upregulation of innate immunity genes in *nuo-6* mutants, we compared gene expression between *nuo-6* and *nuo-6;atfs-1* mutants with wild-type worms and *atfs-1* mutants as controls. Deletion of *atfs-1* significantly decreased the expression of genes involved in innate immunity in *nuo-6* worms, but did not decrease the expression of these genes in wild-type worms. Gene expression changes were determined by RNA sequencing of six biological replicates of each genotype. Results represent counts per million (CPM) expressed as a percentage of wild-type. Worms in a wild-type background (control) are shown with white bars, while worms with *atfs-1* deletion are shown with blue bars. Error bars indicate SEM. **p<0.01, ***p<0.001.

To determine if ATFS-1 is also responsible for the upregulation of innate immunity genes in *nuo-6* mutants, we examined the effect of *atfs-1* deletion. While deletion of *atfs-1* did not decrease the expression of any of the innate immunity genes examined in wild-type worms, loss of *atfs-1* markedly decreased the expression of innate immunity genes in *nuo-6* worms, in most cases reverting expression levels to wild-type (**Fig. 6B**). Thus, ATFS-1 is required for the upregulation of innate immunity genes in *nuo-6* mutants, but is dispensable for the normal nutrient-driven regulation of innate immune gene expression that is seen in wild-type worms(39).

### Activation of ATFS-1 is sufficient to upregulate innate immune signaling genes

Since loss of *atfs-1* prevented the upregulation of innate immunity genes in *nuo-6* mutants, we next sought to determine if activation of ATFS-1 is sufficient to cause upregulation of innate immunity genes. To do this, we examined gene expression in two constitutively active *atfs-1* mutants, *et15* and *et17*(49). We found constitutive activation of ATFS-1 resulted in a significant upregulation of innate immunity genes (**Fig. 7A**). As a positive control, we found that the ATFS-1 target gene *hsp-6* is significantly upregulated in both constitutively activated *atfs-1* mutants (**Fig. S12**).

**Figure. 7.**
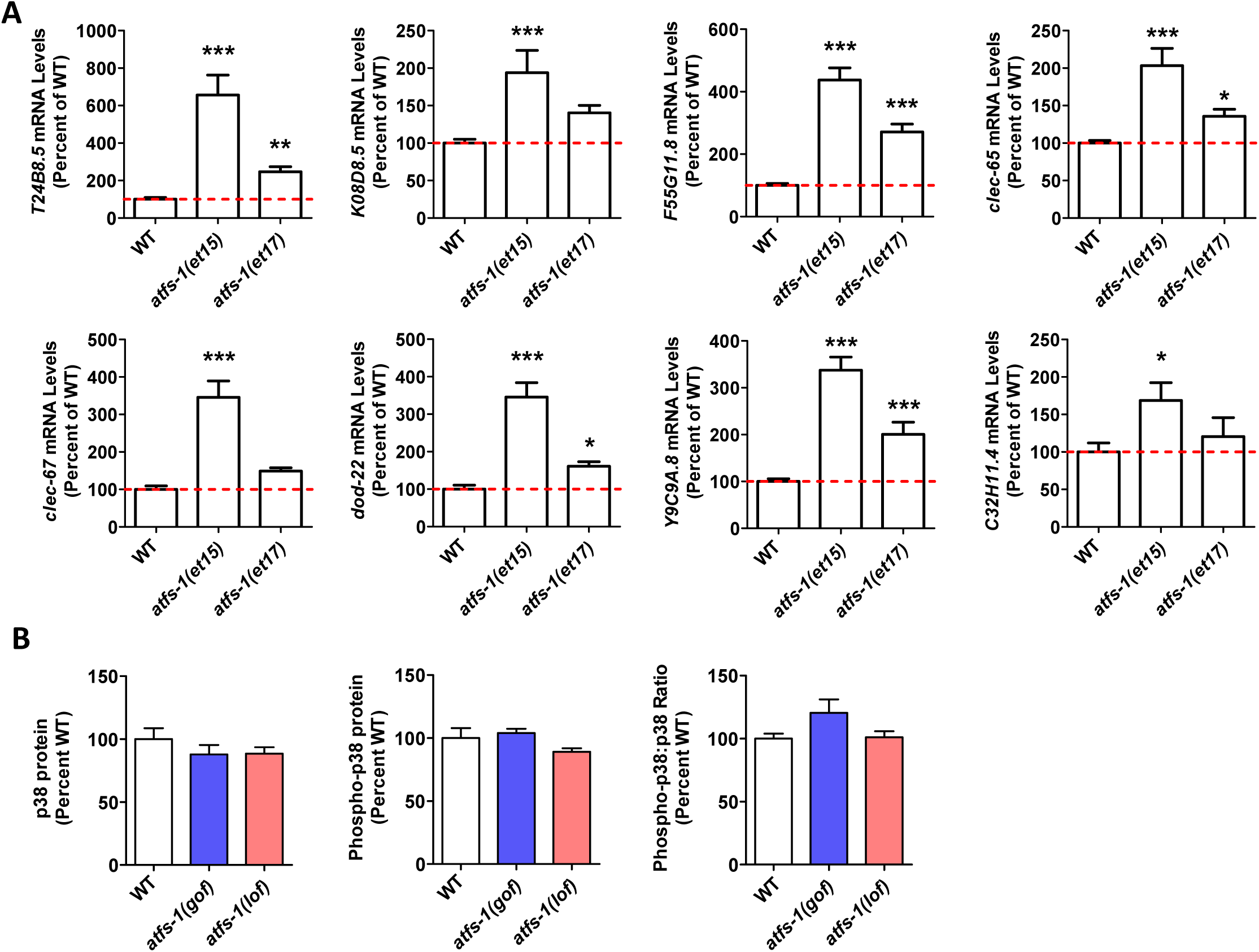
Constitutive activation of mitochondrial unfolded protein response results in upregulation of innate immunity genes. **A.** Examination of mRNA levels of genes involved in innate immunity in two constitutively active *atfs-1* mutants (*et15* and *et17)* revealed that innate immunity genes are upregulated when ATFS-1 is activated. This indicates that activation of the mitochondrial unfolded protein response can cause activation of genes that are regulated by the p38-mediated innate immune pathway. Gene expression changes were determined by RNA sequencing of six biological replicates of each genotype. Results represent counts per million (CPM) expressed as a percentage of wild-type. **B.** Constitutive activation of ATFS-1 does not increase activation of PMK-1/p38 as measured by the ratio of phospho-p38 to total p38 using Western blotting. *atfs-1(gof)* is *et17* allele*. atfs-1(lof)* is *gk3094* allele. Protein levels were quantified for four biological replicates. See **Supplemental Data** for complete images of Western blots including loading controls. Error bars indicate SEM. *p<0.05 **p<0.01, ***p<0.001.

**Figure 8.**
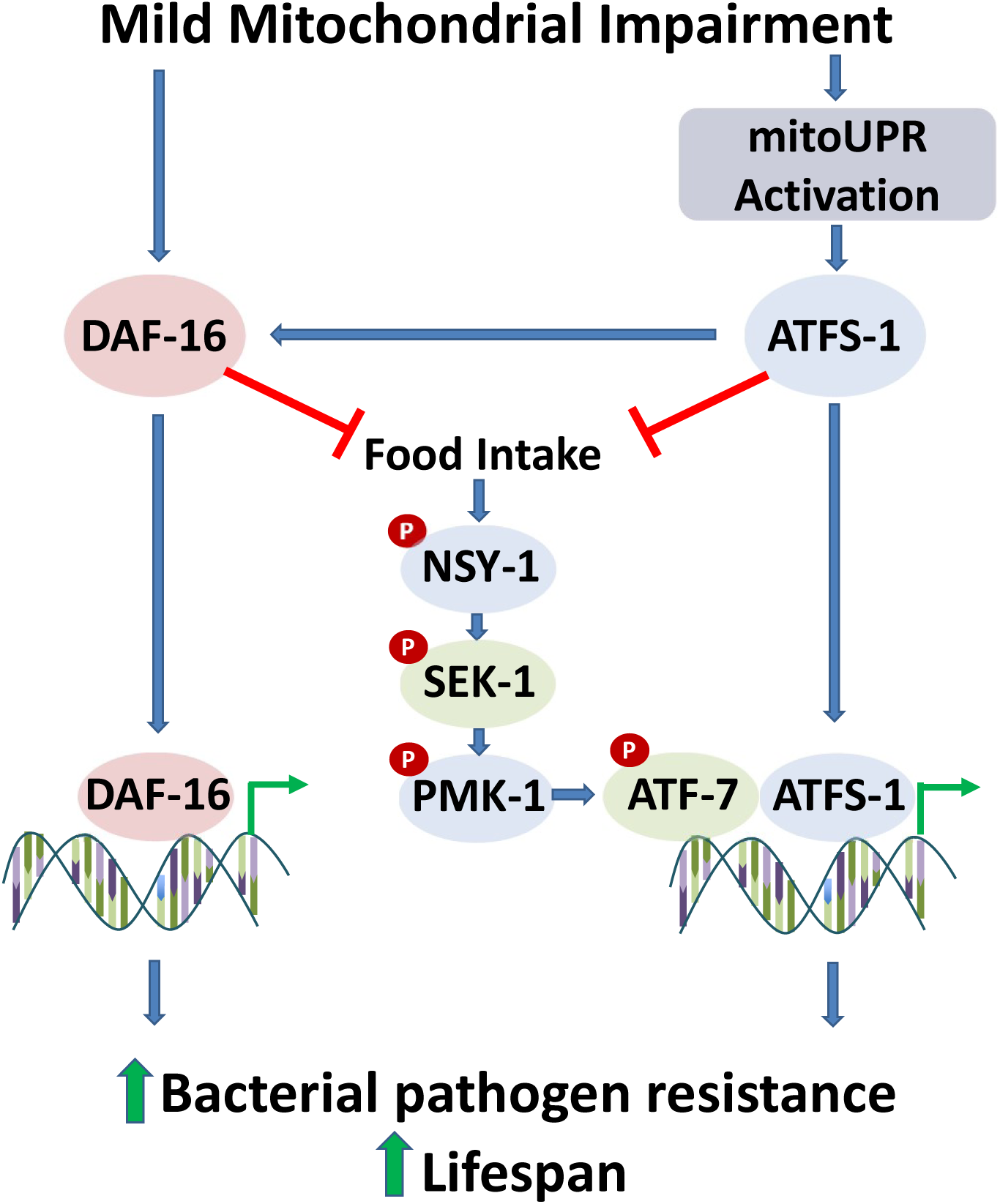
Mild impairment of mitochondrial function increases bacterial pathogen resistance and lifespan. Impairment of mitochondrial function through mutation of *nuo-6* or *isp-1*, which affect the mitochondrial electron transport chain, leads to the activation of the mitochondrial unfolded protein response (mitoUPR) and nuclear localization of DAF-16. The mitoUPR transcription factor ATFS-1 can bind directly to innate immunity genes to increase their expression. Activation of ATFS-1 is also sufficient to decrease food consumption, which results in decreased signaling through the p38-mediated innate immune signaling pathway. This signaling pathway terminates with the ATF-7 transcription factor that can bind to the same genes as ATFS-1. ATFS-1 also acts to facilitate the nuclear localization of DAF-16, which modulates the expression of a separate set of genes that promotes survival of bacterial pathogens and longevity. DAF-16 acts to decrease food intake thereby decreasing signaling through the p38-mediated innate immune signaling pathway. DAF-16, ATFS-1, and all of the components of the p38-mediated innate immune signaling pathway (NSY-1, SEK-1, PMK-1 and ATF-7) are required for the increased resistance to bacterial pathogens and increased lifespan in *nuo-6* and *isp-1* mutants.

To determine if activation of ATFS-1 causes upregulation of innate immunity genes through activation of the p38-mediated innate immune signaling pathway, we examined PMK-1/p38 phosphorylation in a constitutively activated *atfs-1* mutant (*et17*) and an *atfs-1* loss-of-function mutant (*gk3094*). Similar to what we observed in the long-lived mitochondrial mutants, we found that the levels of PMK-1/p38 protein, phosphorylated PMK-1/p38 protein and the ratio of phosphorylated p38/p38 are equivalent to wild-type in both *atfs-1* mutants (**Fig. 7B**). This suggests that activation of ATFS-1 can upregulate innate immunity genes without increased activation of the p38-mediated innate immune signaling pathway.

To determine if ATFS-1 might be able to directly bind to the same innate immunity genes as ATF-7, we compared data from two previous CHiP-seq studies(36, 50). For this comparison, we defined innate immunity genes as genes that are upregulated in response to a 4-hour exposure to PA14(36). After exposure to PA14, ATF-7 was found to be bound to 345 of these innate immunity genes(36). Following exposure to mitochondrial stress induced by *spg-7* RNAi, ATFS-1 was found to be bound to 51 of these innate immunity genes(50). Of the 51 innate immunity genes that can be bound by ATFS-1, 49% (25 genes) can also be bound by ATF-7 (**Fig. S13; Table S3**). This binding likely occurs at different sites since the predicted binding site motifs of ATFS-1 and ATF-7 bear little similarity(36, 50). The fact that ATFS-1 and ATF-7 are able to bind to the same genes involved in innate immunity indicates that the mitoUPR and p38-mediated innate immune signaling pathway act in parallel to modulate the expression of innate immunity genes, which are critical for lifespan extension.

## Discussion

Aging may be defined as a progressive decline of physiologic functions that increases the probability of death. The major factors that contribute to aging have been categorized into hallmarks or pillars of aging(51). Interestingly, at least two of the seven pillars are directly related to the immune system, but in opposite directions. First, the ability of the immune system to fight off external stresses such as exposure to bacterial pathogens declines with age, which can be accounted for by an age-dependent decrease in basal and activated PMK-1/p38 levels(52, 53). Second, there is an increase in the basal activity of the immune system that occurs with advancing age, eventually leading to chronic inflammation(54). Thus, the relationship between aging and the immune response can be seen as a fine balance between maintaining the ability to respond to stresses, while limiting non-specific immune activation.

Our ability to precisely define the relationship between immune activity and aging has been greatly enhanced by the discovery of genes that increase lifespan in model organisms, which allow for investigation of the molecular mechanisms. In this work, we demonstrate that mild impairment of mitochondrial function through mutations affecting either complex I (*nuo-6*) or complex III (*isp-1*) of the electron transport chain results in increased expression of a broad panel of genes involved in innate immunity. Treatment with the complex I inhibitor rotenone was previously shown to cause neurodegeneration and the upregulation of a small panel of genes involved in innate immunity, but the mechanism by which these genes were upregulated was not determined(37). While it is well established that the degeneration of neurons can activate an innate immune response (55, 56), our work clearly demonstrates that the impairment of mitochondrial function can also lead to the upregulation of innate immunity genes. Further, our results indicate that the upregulation of innate immunity genes in response to mild mitochondrial impairment is dependent on the mitoUPR transcription factor ATFS-1 and all of the components of the p38-mediated innate immune signaling pathway (NSY-1, SEK-1, PMK-1, ATF-7).

While both the long-lived mitochondrial mutants and constitutively active *atfs-1* mutants, exhibit increased expression of ATF-7 target genes overall, these mutants show no increase in p38-mediated innate immune signaling (**Fig. 7**). Moreover, since ATFS-1 activation is sufficient to decrease food intake, these mutants would be predicted to have decreased ATF-7 target gene expression because decreasing food intake diminishes p38-mediated innate immune signaling(39). Combined this suggests that ATFS-1 can activate innate immunity genes directly. This conclusion is supported by the fact that ATFS-1 can bind to the same innate immunity genes as ATF-7 (**Fig. S12**).

Previous work examining the relationship between the mitoUPR and p38-mediated innate immune signaling pathway has generated conflicting results. In an earlier study, activation of the mitoUPR by *spg-7* RNAi was found to increase the survival of *pmk-1* and *sek-1* mutants on PA14, leading the authors to conclude that the mitoUPR was increasing pathogen resistance independently of the p38-mediated innate immune signaling pathway(29). Subsequently, it was shown that overexpression of the mitoUPR target gene *hsp-60* is sufficient to activate the p38-mediated innate immune signaling pathway leading to increased expression of genes involved in innate immunity and increased PA14 resistance, all of which was dependent on p38-mediated innate immune signaling pathway (*nsy-1, sek-1* and *pmk-1*)(40). Our current results resolve this discrepancy by demonstrating that the mitoUPR can affect the expression of innate immunity genes and pathogen resistance both directly through binding of ATFS-1 to innate immunity genes, and indirectly through the p38-mediated innate immune signaling and ATF-7 (**Fig. 7**).

The increased expression of innate immunity genes in *nuo-6* and *isp-1* worms results in enhanced resistance to bacterial pathogens, which is dependent on both ATFS-1 and the p38-mediated innate immune signaling pathway. The increased resistance to bacterial pathogens is also dependent on DAF-16, which has increased nuclear localization in long-lived mitochondrial mutants(20). However, the mechanism by which DAF-16 protects against bacterial pathogens appears to be distinct from the mechanism of ATFS-1, as loss of DAF-16 does not affect the expression of the innate immunity genes that are upregulated by ATFS-1 and ATF-7 (**Fig. 5; Fig. S10**)(38). The effect of DAF-16 on bacterial pathogen resistance may instead be due to a general effect of DAF-16 on resistance to stress.

We have previously shown that both ATFS-1 and DAF-16 are required for the long-lifespan of *nuo-6* and *isp-1* mutants(19, 20). Here, we show that their lifespan is also completely dependent on the p38-mediated innate immune signaling pathway (NSY-1, SEK-1, PMK-1, ATF-7). As all of these factors are also required for the enhanced bacterial pathogen resistance in *nuo-6* and *isp-1* worms, this indicates that there are multiple conserved genetic pathways acting in parallel to upregulate innate immunity and promote longevity in response to impaired mitochondrial function.

It appears that there are multiple ways to optimize innate immune signaling with respect to longevity. As in the long-lived mitochondrial mutants that we studied here, the p38-mediated innate immune signaling pathway is also required for the longevity of other genes and interventions that increase lifespan. *daf-2* mutants have increased lifespan(1) and increased resistance to bacterial pathogens(57). Unlike the *nuo-6* and *isp-1* mutants, *daf-2* mutants show a downregulation of many, but not all, SEK-1-dependent genes involved in innate immunity through decreased feeding and decreased phosphorylation of PMK-1/p38(39). Nonetheless, genes involved in the p38-mediated innate immune signaling pathway (SEK-1, PMK-1) are required for the long-lifespan, and increased resistance to bacterial pathogens in *daf-2* worms, as is DAF-16(28, 38, 58).

Similar to *daf-2* mutants, the increase in lifespan caused by dietary restriction resulting from decreased nutrient availability is dependent on the p38-mediated innate immune signaling pathway (NSY-1, SEK-1, PMK-1, ATFS-7), despite the fact that dietary restriction was found to decrease PMK-1/p38 activation and downregulate more innate immune genes (162 genes) than were upregulated (46 genes)(39). Based on this and other supporting data it was concluded that lifespan extension resulting from dietary restriction was not mediated by pathogen resistance per se, but by a metabolic/nutrient-related function of the p38-mediated innate immune signaling pathway.

Combined, this clearly indicates that it is not a simple relationship between innate immune activation and maximum longevity, as increased lifespan can be achieved with upregulation or downregulation of innate immunity genes. While a functional p38-mediated innate immune signaling pathway may be universally required for long life, different levels of its activity are required to achieve maximum lifespan extension under different circumstances or genetic backgrounds. While many previous studies have only examined a few genes involved in innate immunity, the fact that different innate immune genes can be both upregulated and downregulated in the same mutants demonstrates the importance of looking more broadly across all genes involved in innate immunity to gain a more in-depth understanding of how these changes are affecting longevity. Moreover, it will be important to understand the specific roles of each of these genes in pathogen defense and longevity.

## Conclusions

Overall, this work demonstrates the importance of the p38-mediated innate immune signaling pathway and the mitochondrial unfolded protein response for both pathogen resistance and lifespan. It shows that activation of innate immunity is a key function of the mitoUPR that works in concert with the p38-mediated innate immune signaling pathway as a key defense mechanism, which is activated by mitochondrial impairment. The relationship between innate immune gene expression, pathogen resistance and lifespan is complex, and there are multiple ways to optimize the levels of innate immune activation with respect to longevity. As all of the signaling pathways studied here are conserved up to mammals, advancing our understanding of the links between innate immune gene expression, pathogen resistance and longevity in model organisms, may advance our understanding of these processes in humans. Understanding the relationship between mitochondrial function, innate immunity and longevity will provide important insights into fundamental aspects of how lifespan can be extended.

## Materials and Methods

### Strains

WT(N2)

MQ1333 *nuo-6(qm200)*

JVR171 *isp-1(qm150)*

T24B8.5p::GFP

JVR521 *nsy-1(ok593)*

JVR520 *sek-1(km4)*

JVR165 *pmk-1(km25)*

ZD442 *atf-7(qd22); agIs219[T24B8.5p::GFP+ttx-3p::GFP]*

JVR455 *atfs-1(gk3094)*

JVR479 *nuo-6(qm200);atfs-1(gk3094)*

QC1115 *atfs-1(et15)*

QC1117 *atfs-1(et17)*

JVR533 *nuo-6(qm200);nsy-1(ok593)*

JVR543 *nuo-6(qm200);sek-1(km4)*

JVR281 *nuo-6(qm200);pmk-1(km25)*

JVR527 *nuo-6(qm200);atf-7(qd22); agIs219[T24B8.5p::GFP+ttx-3p::GFP]*

JVR532 *isp-1(qm150);nsy-1(ok593)*

JVR534 *isp-1(qm150);sek-1(km4)*

JVR538 *isp-1(qm150);atf-7(qd22); agIs219[T24B8.5p::GFP+ttx-3p::GFP]*

*math-33(tm6724)*

JVR456 *isp-1(qm150); math-33(tm6724)*

JVR457 *nuo-6(qm200); math-33(tm6724)*

*daf-16(mu86)*

JVR380 *isp-1(qm150);daf-16(mu86)*

Strains were grown on OP50 bacteria at 20°C. All of the strains were genotyped or sequenced to confirm the presence of the mutation in the gene of interest. Double mutants were generated by crossing WT males to *nuo-6* or *isp-1* mutants and crossing the resulting heterozygous male progeny (*nuo-6/+* or *isp-1/+*) to the mutant of interest (*nsy-1, sek-1, pmk-1, atf-7*). After selfing, slow growing progeny were singled and genotyped or sequenced to identify double mutants. We were unable to generate *isp-1 pmk-1* and *nuo-6 daf-16* double mutants due to the close proximity of these two genes on the same chromosome.

### Bacteria pathogenesis – slow kill assay

Pathogenesis assays with *P. aeruginosa* strain PA14 were performed as described previously(39). In brief, the overnight PA14 culture was seeded to the center of a 35-mm NGM agar plate containing 20 mg/L FUdR. Seeded plates were incubated at 37°C overnight, then incubated at room temperature overnight. Approximately 40 day three adults were transferred to these prepared plates, with 10 plates scored per strain. The assays were conducted at 20°C. Animals that did not respond to gentle prodding from a platinum wire were scored as dead.

### Lifespan

All lifespan assays were performed at 20°C. Except where noted lifespan assays were performed on NGM plates seeded with OP50 bacteria. Lifespan assays included FUdR to limit the development of progeny. Lifespan assays on solid plates utilized 25 µM FUdR to minimize potential effects of FUdR on lifespan(59). Animals were excluded from the experiment if they crawled off the plate or died of internal hatching of progeny or expulsion of internal organs. *Lifespan assays with arrested bacteria.* Growth-arrested bacterial food was prepared through treatment with antibiotics and cold as described previously(39). All worms were passaged at 20°C for at least two generations before lifespan assays were initiated. Synchronized populations of L1 animals were obtained by hypochlorite treatment, then allowed to develop at 20°C on standard NGM plates seeded with OP50-1. The young adults were transferred to NGM agar plates containing 50 mg/L ampicillin, 10 mg/L kanamycin, 1 mg/L tetracycline, 50mg/L nystatin, and 100 mg/L FUdR, and seeded with the cold- and antibiotic-treated OP50-1. Three days later the animals were washed twice in S basal, and transferred to a 24 well plate with 1 mL S basal in each well containing the arrested OP50-1 at OD_600_ 3. 20–40 worms were placed in each well.

### Western Blotting

Protein levels were measured by western blotting in whole worm lysates obtained from four independent biological replicates. Briefly, whole worm lysates obtained from approximately 1000 animals on day 1 of adulthood were subjected to SDS-PAGE in polyacrylamide gels (4%-12%), immunoblotted with specific antibodies against phospho-p38 (Cell Signaling) or total p38/PMK-1(47), and visualized following standard procedures. Quantification analysis of all blots was performed with the use of ImageJ software. Results were corrected to Ponceau red staining (0.5%, w:v) of the membrane since *nuo-6* worms were found to exhibit decreased levels of tubulin protein.

### Food intake assay

Food intake was quantified in liquid culture by measuring the relative amount of growth-arrested bacteria that are present in a culture before and after incubation with *C. elegans*, as previously described(39). Synchronized populations of L1 animals were obtained by hypochlorite treatment and allowed to develop at 20°C on standard NGM agar plates seeded with OP50. The worms were transferred onto NGM plates containing FUdR at the L4 stage of development. On day 3 of adulthood, animals were washed twice in S basal (5.85 g/L NaCl, 1 g/L K2HPO4, 6 g/L KH2PO4) and 30-50 worms were transferred to each well of a 24 well plate containing 1 mL of arrested OP50 at optical density 3 (600 nm) re-suspended in S basal complete medium (S basal containing 5 mg/L cholesterol, 50 mg/L ampicillin, 10 mg/L kanamycin, 1 mg/L tetracycline, 50 mg/L nystatin, FUdR). On day 12 of adulthood, bacteria were removed from the wells by washing and diluted 10x with S basal prior to measuring the optical density (600 nm). The relative food intake was determined by the change in OP50 optical density between day 3 and day 12 of adulthood, and normalized to the number of worms per well.

### Bacterial avoidance assay

A colony of PA14 was placed in 5 ml of Luria Broth (LB) or a colony of OP50 was placed in 5ml 2YT, and the cultures were incubated, shaking, for 16 hr at 37*°*C. 100 ul of the culture was seeded onto the center of 6 cm NGM plates, which were grown for 24 hours at room temperature. Animals were synchronized by picking stage L4 animals the day before the experiment and were placed on OP50 plates at 20*°*C. The synchronized day 1 animals were washed off of the OP50 plates with M9 buffer, washed three times in M9 buffer, and then ∼40 animals were deposited on each of the assay plates using a glass pipette. The animals were placed on the agar ∼1 cm from the edge of the bacterial spots. The plates were then placed at 20*°*C. The number of animals on and off of the bacteria were counted after 24 hours.

### Quantification of reporter fluorescence

*T24B8.5p::GFP* animals were randomly selected and imaged with ZEN 2012 software on an Axio Imager M2 microscope with a 10X/0.25 objective (Zeiss, Jena, Germany). Fluorescence brightness was quantified blindly using NIH ImageJ software.

### RNA isolation

mRNA was collected from pre-fertile young adult worms using Trizol as previously described (60). For quantitative real-time RT-PCR, we collected three biological replicates for each strain on separate days. For RNA sequencing experiments, we collected 3-6 biological replicates.

### Quantitative Real-Time RT-PCR

The mRNA was converted to cDNA using a High-Capacity cDNA Reverse Transcription kit (Life Technologies/Invitrogen) according to the manufacturer’s directions. qPCR was performed using a FastStart Universal SYBR Green kit (Roche) in a Bio-Rad iCycler real-time PCR detection system (61). Primer sequences for other genes tested include:

T24B8.5 (TACACTGCTTCAGAGTCGTG, CGACAACCACTTCTAACATCTG);

K08D8.5 (CAAAATATCCTCCGGGAAGTC, TTCACGGAATCACCATCGTA);

F55G11.8 (GGAAATGGTTGCAAACTTGG, TGCAGAATCGACAGTTTGGA);

*clec-67* (TTTGGCAGTCTACGCTCGTT, CTCCTGGTGTGTCCCATTTT);

*clec-65* (GCAATCAACCTCGTGATGTG, CGCAGAAGCAGTTTGTATCC);

*dod-22* (TCCAGGATACAGAATACGTACAAGA, GCCGTTGATAGTTTCGGTGT);

Y9C9A.8 (CGGGGATATAACTGATAGAATGG, CAAACTCTCCAGCTTCCAACA); and

C32H11.4 (GGCAATACTTGCAAGACTGGAA, CCAGCTACACGATTGGTCCT).

### RNA sequencing and Bioinformatic Analysis

RNA sequencing was performed as previously described(62). RNA-seq data is available on NCBI GEO: GSE93724(20), GSE110984(19) and was analyzed by the Harvard School of Public Health Bioinformatics core for this paper.

#### Read mapping and expression level estimation

All samples were processed using an RNA-seq pipeline implemented in the bcbio-nextgen project (https://bcbio-nextgen.readthedocs.org/en/latest/). Raw reads were examined for quality issues using FastQC (http://www.bioinformatics.babraham.ac.uk/projects/fastqc/) to ensure library generation and sequencing data were suitable for further analysis. If necessary, adapter sequences, other contaminant sequences such as polyA tails and low quality sequences were trimmed from reads using cutadapt http://code.google.com/p/cutadapt/. Trimmed reads were aligned to the Ensembl build WBcel235 (release 90) of the *C.elegans* genome using STAR(63). Quality of alignments was assessed by checking for evenness of coverage, rRNA content, genomic context of alignments (for example, alignments in known transcripts and introns), complexity and other quality checks. Expression quantification was performed with Salmon(64) to identify transcript-level abundance estimates and then collapsed down to the gene-level using the R Bioconductor package tximport(65). Principal components analysis (PCA) and hierarchical clustering methods validated clustering of samples from the same batches and across different mutants.

#### Differential gene expression and functional enrichment analysis

Differential expression was performed at the gene level using the R Bioconductor package DESeq2(66). For each wildtype-mutant comparison significant genes were identified using an FDR threshold of 0.01. For datasets in which experiments were run across two batches, we included batch as a covariate in the linear model.

#### Overlap table

Lists of differentially expressed genes were separated by direction of expression change and compared to genes that are modulated in response to exposure to the bacterial pathogen *P. aeruginosa* strain PA14 in a PMK-1 and ATF-7-dependent manner, as identified by Fletcher et al., 2019(36). Significance of overlap was computed using the hypergeometric test. *Heatmaps.* For each dataset (long-lived mutant comparison to wildtype), the raw counts were regularized log (rlog) transformed using the DESeq2 package(66). This transformation moderates the variance across the mean, improving the clustering in an unbiased manner. The rlog matrix was then subset to retain only those genes from the Fletcher et al gene lists which were also significantly differentially expressed between mutant and wildtype. Heatmaps were plotted using the pheatmap package in R.

### Statistical Analysis

Experiments were conducted such that the experimenter was blinded to the genotype of the animals being tested. The worms for each experiment were randomly selected from maintenance plates at stages when all worms appeared healthy. Except where noted, we completed a minimum of three biological replicates for each assay (independent population of worms tested on a different day). Statistically significance of differences between groups of more than two was determined by one-way ANOVA with Dunnett’s multiple comparison post-hoc test or two-way ANOVA with Bonferroni post-doc test using Graphpad Prism. For survival analysis, a log-rank test was used. On all bar graphs, the bar indicates the average (mean) and error bars indicate standard error of the mean (SEM). This study was not pre-registered. No sample size calculations were performed. This study did not include a pre-specified primary endpoint.

## Acknowledgments

Some strains were provided by the CGC, which is funded by NIH Office of Research Infrastructure Programs (P30 OD010440). We would also like to acknowledge the *C. elegans* knockout consortium and the National Bioresource Project of Japan for providing strains used in this research.

## Author Contributions

Conceptualization: JVR, TKB. Methodology: JCC, ZW, PR, SS, MM, TKB, JVR. Investigation: JCC, ZW, PR, SS, MM, JVR. Visualization: JCC, ZW, PR, SS, MM, JVR. Writing – original draft: JVR. Writing – review and editing: JCC, ZW, PR, SS, MM, TKB, JVR. Supervision: JCBF, TKB, JVR.

## Materials & Correspondence

Correspondence and material requests should be addressed to Jeremy Van Raamsdonk.

## Data availability

RNA-seq data has been deposited on GEO: GSE93724, GSE110984. All other data and strains generated in the current study are included with the manuscript or available from the corresponding author on request.

## Supplementary Figures

This PDF file includes:

Figures S1 to S12

Supplementary data: Western blot image

Other supplementary materials for this manuscript include the following:

Tables S1 to S3

**Figure S1.**
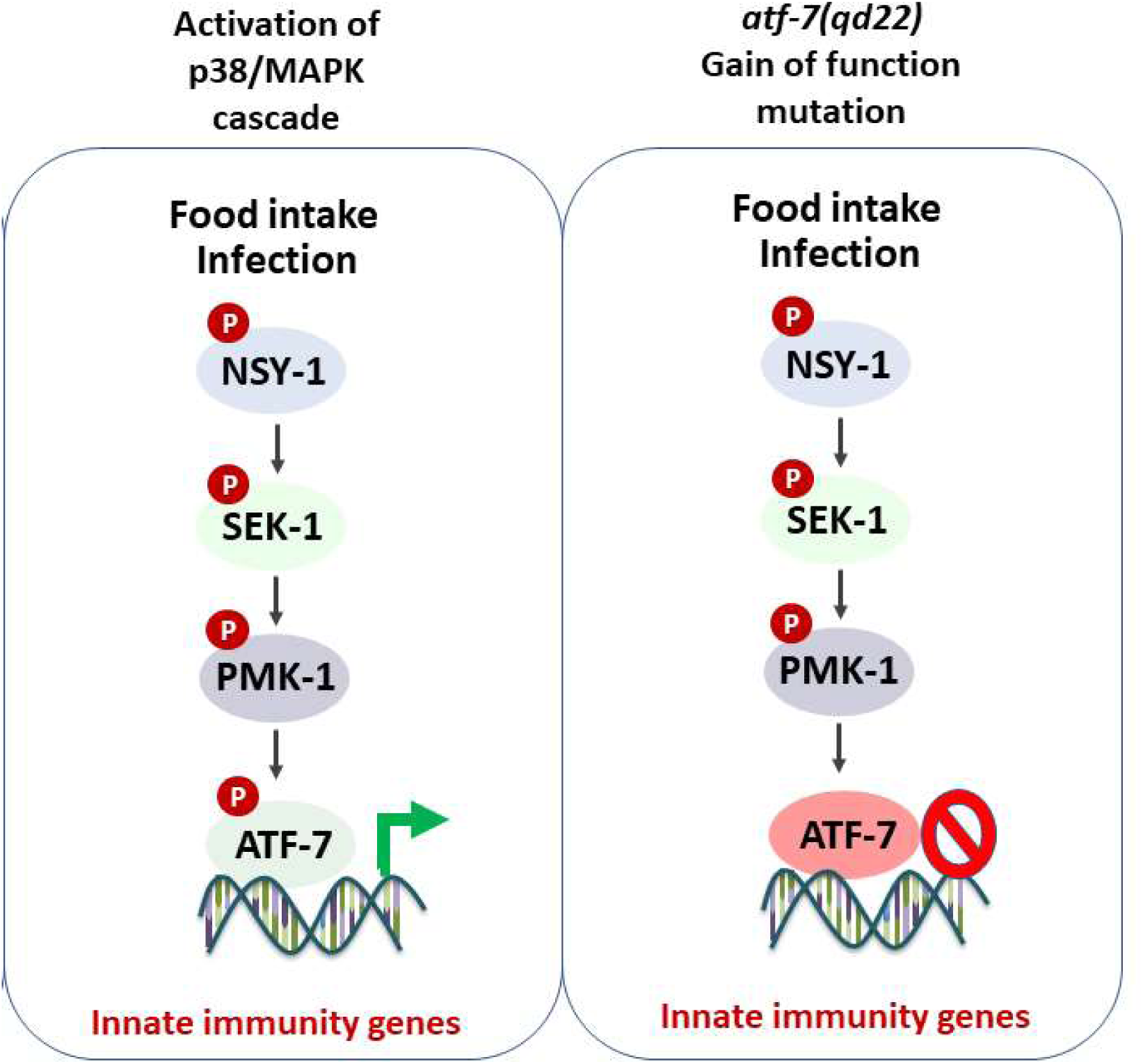
Overview of p38-mediated innate immune signaling pathway. The p38-mediated innate immune signaling pathway is a MAPK signaling pathway. Exposure to bacterial pathogens or increased food intake results in the activation of this pathway through the phosphorylation of NSY-1/ASK1 (MAPK kinase kinase). NSY-1 then phosphorylates SEK-1/MKK3/MKK6 (MAPK kinase), which phosphorylates PMK-1/p38 (MAPK), which phosphorylates the transcription factor ATF-7/ATF2/ATF7/CREB5. Under normal conditions, ATF-7 acts as a repressor inhibiting the expression of innate immunity genes. When ATF-7 is phosphorylated by PMK-1, it becomes an activator promoting the expression of innate immunity genes. The *qd22* mutation prevents phosphorylation of ATF-7 by PMK-1. As a result, ATF-7 with the *qd22* mutation acts as a constitutive repressor even in the presence of bacterial pathogens.

**Figure S2.**
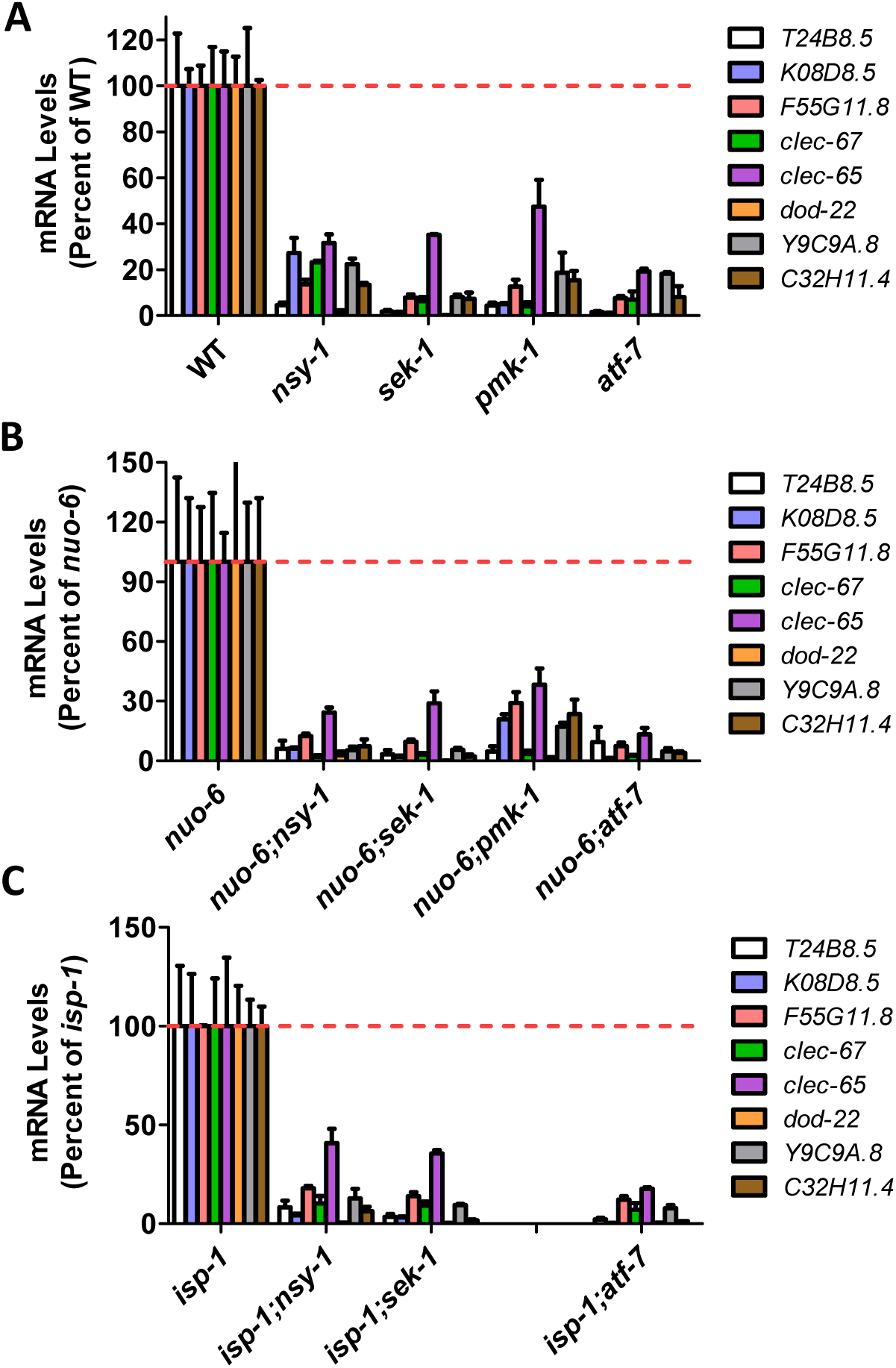
Upregulation of innate immunity genes in long-lived mitochondrial mutants requires the p38-mediated innate immune signaling pathway. Mutation of genes involved in the p38-mediated innate immune signaling pathway (*nsy-1, sek-1, pmk-1, atfs-7(gof)*) decrease the expression of genes involved in innate immunity in wild-type (**A**), *nuo-6* (**B**), and *isp-1* (**C**) worms. Gene expression was determined by quantitative real-time RT-PCR on three biological replicates of pre-fertile young adult worms. Error bars indicate SEM. All differences from control are significant p<0.05.

**Figure S3.**
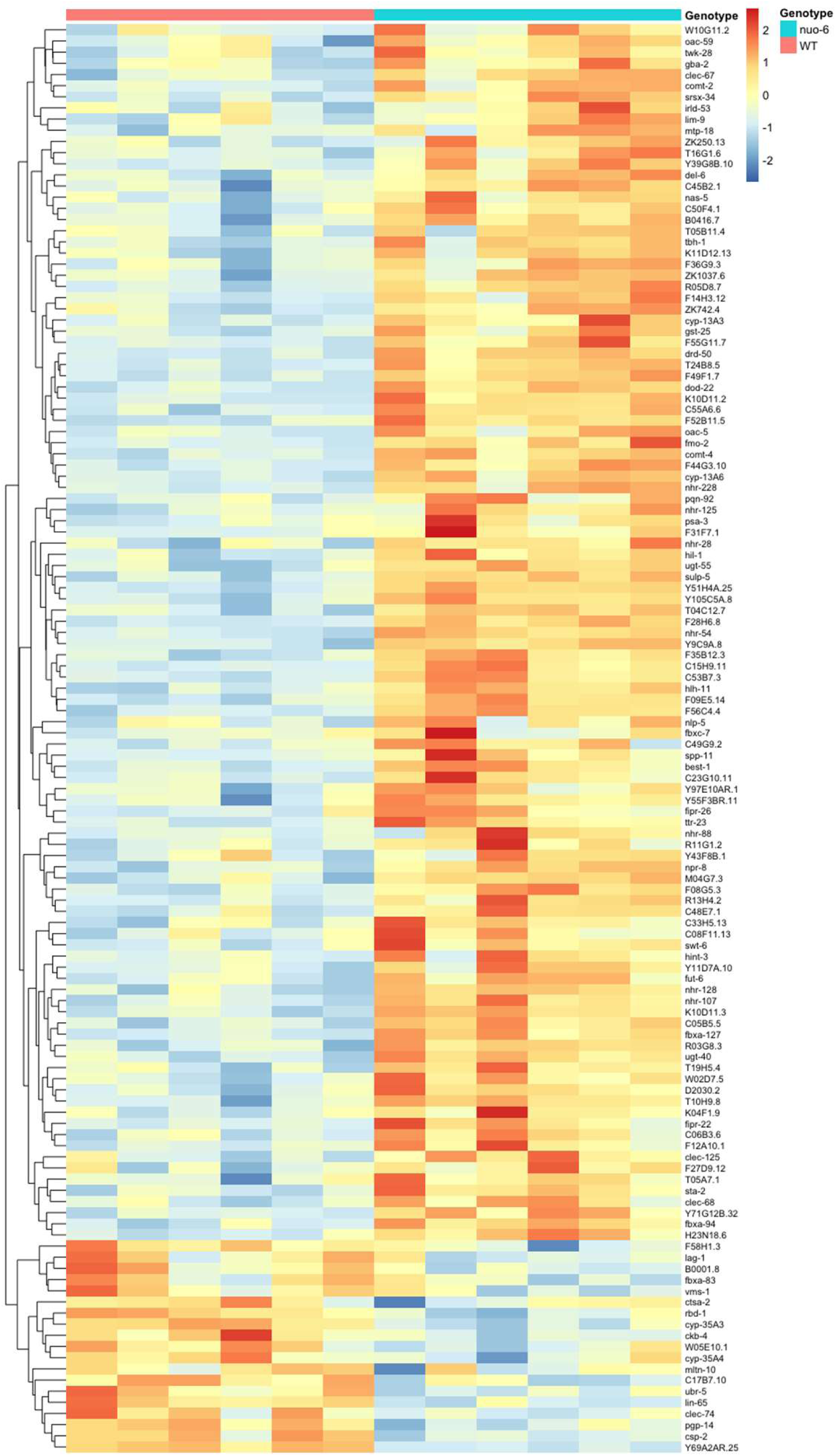
Genes that are upregulated by activation of the p38-mediated innate immune pathway are primarily upregulated in the long-lived mitochondrial mutant *nuo-6*. This heatmap includes genes that are upregulated by exposure to the bacteria pathogen *P. aeruginosa* PA14 in a PMK-1- and ATF-7-dependent manner and for which the expression is significantly changed in *nuo-6* mutants.

**Figure S4.**
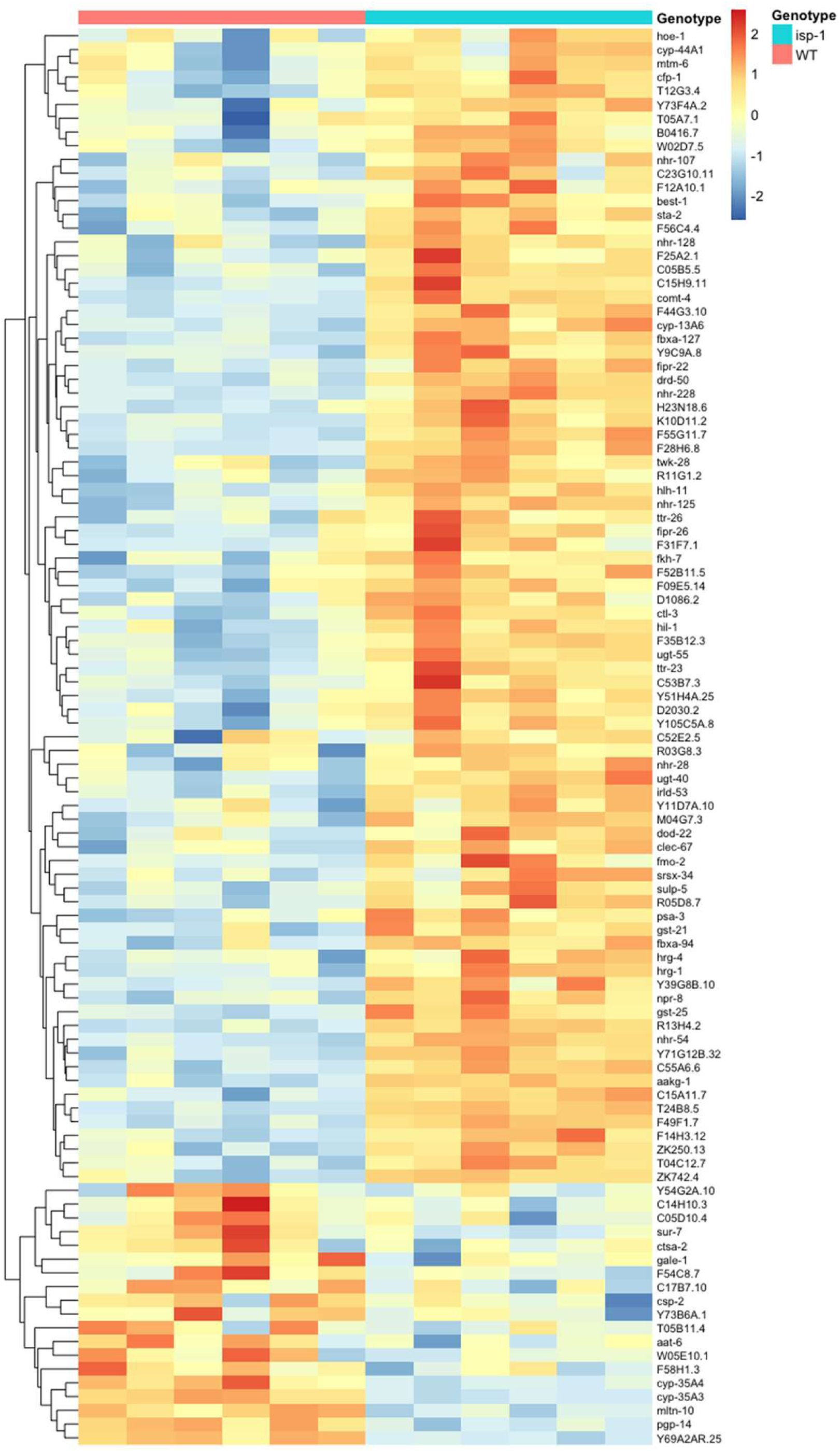
Genes that are upregulated by activation of the p38-mediated innate immune pathway are primarily upregulated in the long-lived mitochondrial mutant *isp-1*. This heatmap includes genes that are upregulated by exposure to the bacteria pathogen *P. aeruginosa* PA14 in a PMK-1- and ATF-7-dependent manner and for which the expression is significantly changed in *isp-1* mutants.

**Figure S5.**
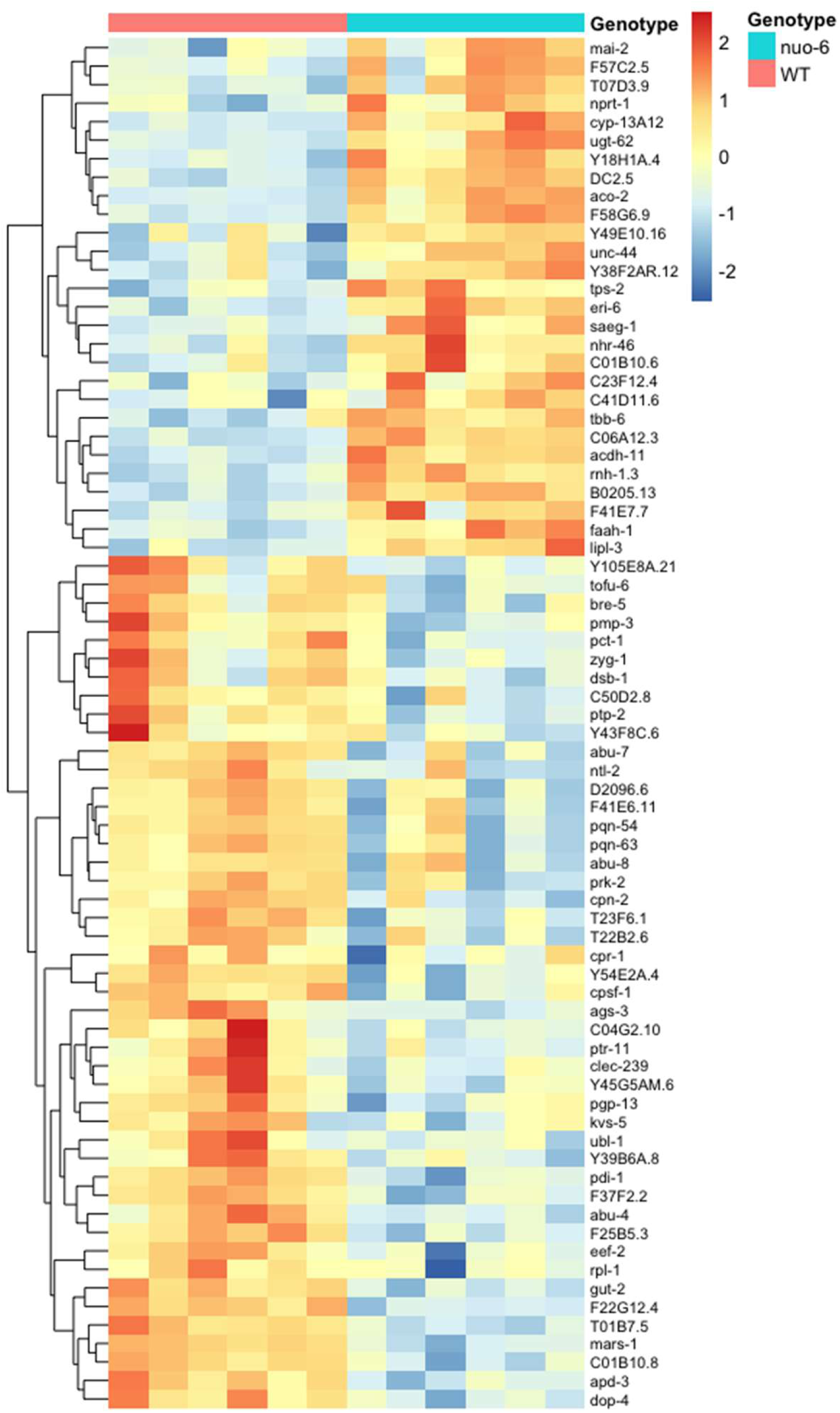
Expression of genes that are downregulated by activation of the p38-mediated innate immune pathway is altered in the long-lived mitochondrial mutant *nuo-6*. This heatmap includes genes that are downregulated by exposure to the bacteria pathogen *P. aeruginosa* PA14 in a PMK-1- and ATF-7-*-*dependent manner and for which the expression is significantly changed in *nuo-6* mutants.

**Figure S6.**
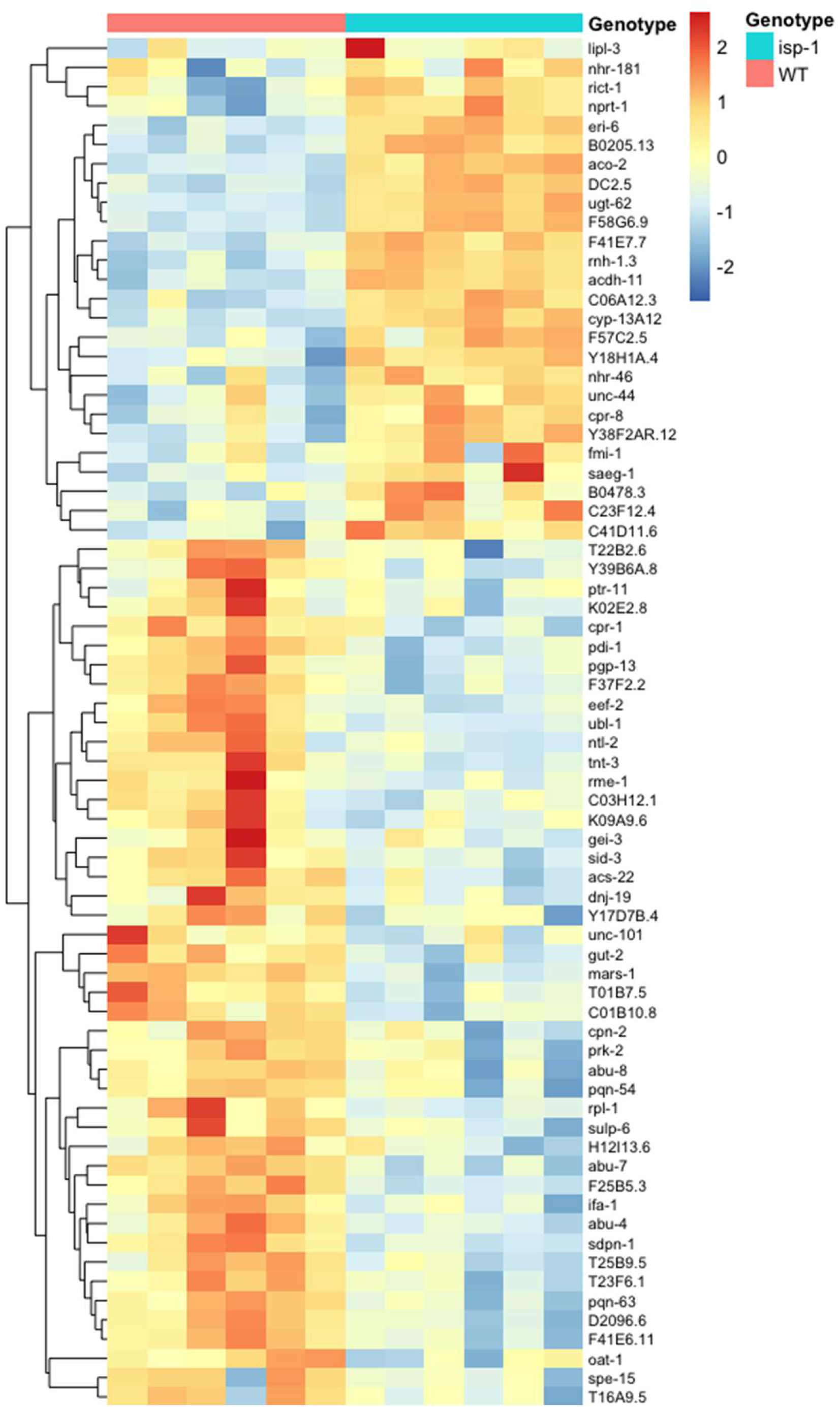
Expression of genes that are downregulated by activation of the p38-mediated innate immune pathway is altered in the long-lived mitochondrial mutant *isp-1*. This heatmap includes genes that are downregulated by exposure to the bacteria pathogen *P. aeruginosa* PA14 in a PMK-1- and ATF-7-*-*dependent manner and for which the expression is significantly changed in *isp-1* mutants.

**Figure S7.**
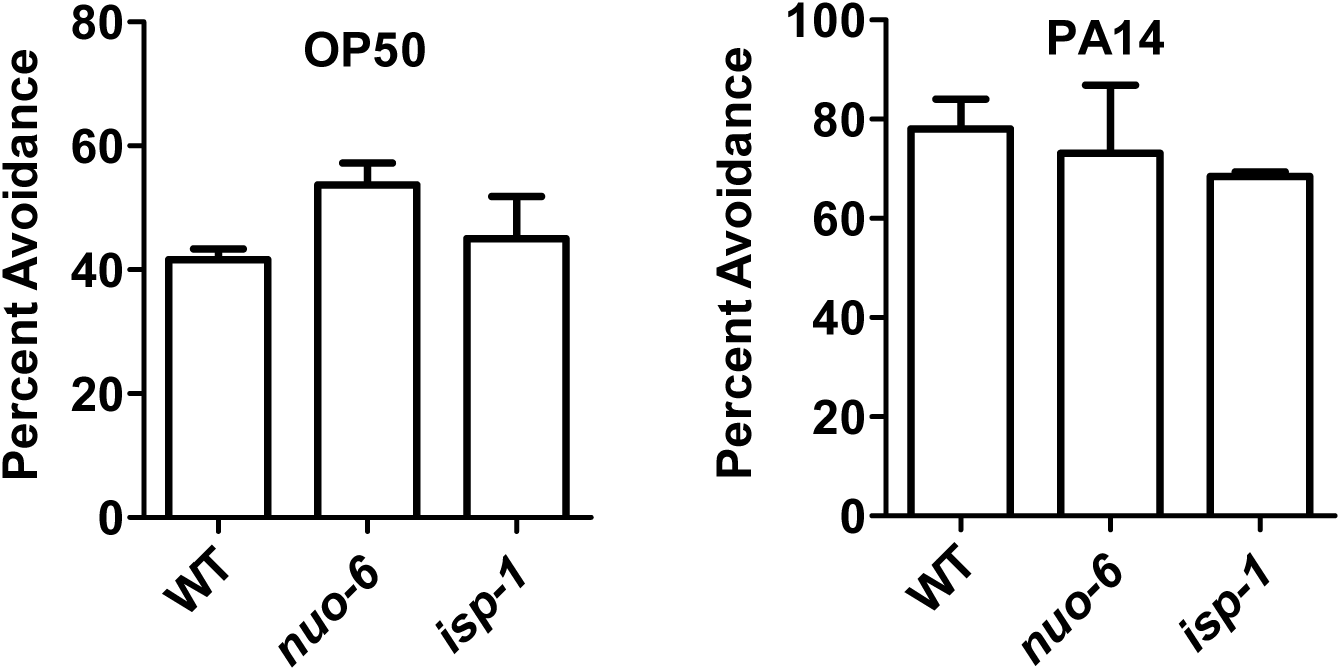
Increased resistance to bacterial pathogens does not result from bacterial avoidance. To explore the mechanisms underlying the increased resistance to bacterial pathogens in *nuo-6* and *isp-1* mutants, bacterial avoidance and food consumption were examined. There was no significant difference in bacterial avoidance between wild-type worms and *nuo-6* or *isp-1* mutants on OP50 bacteria (**A**) or *P. aeruginosa* (**B**). Three biological replicates per strain were quantified. Error bars indicate SEM.

**Figure S8.**
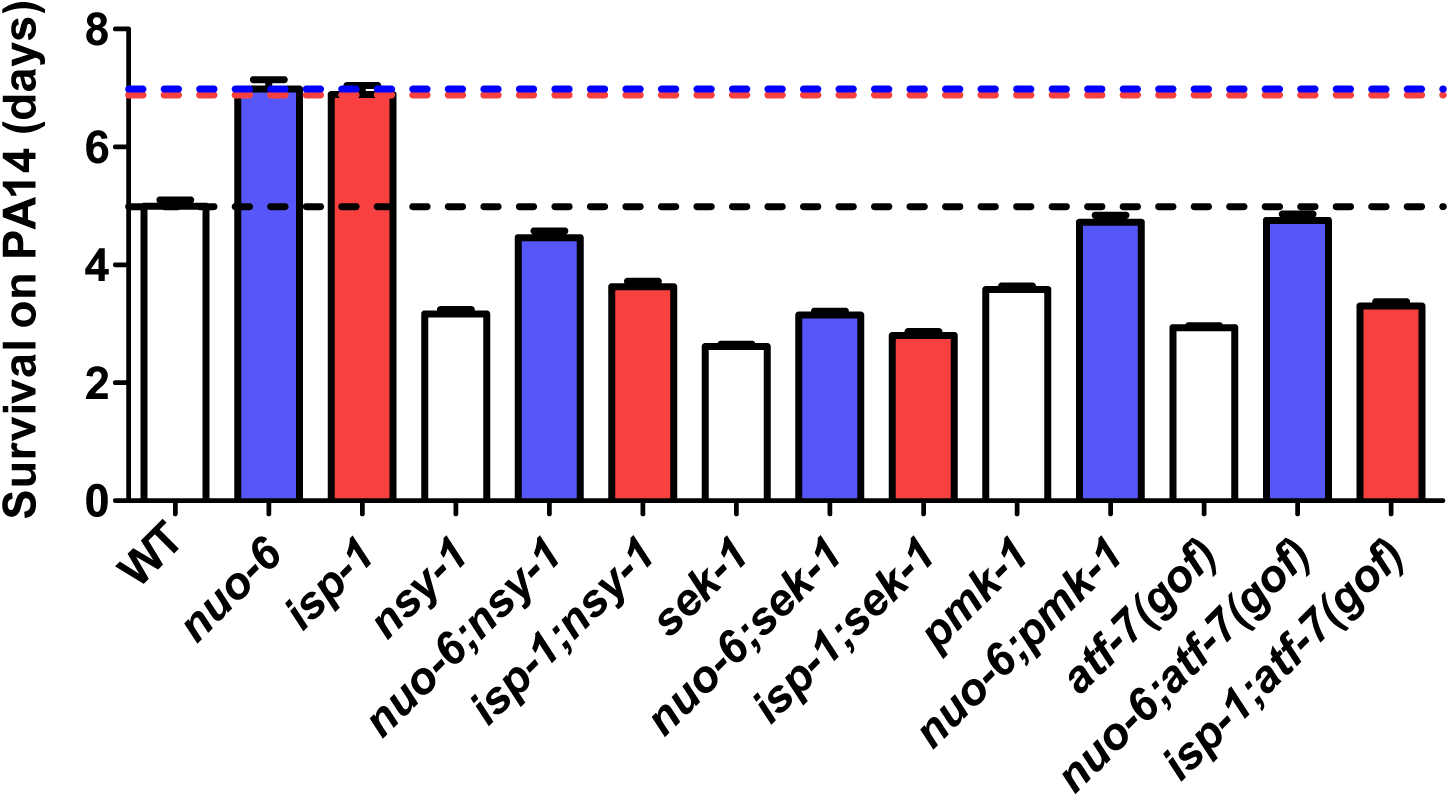
Disruption of p38-mediated innate immune signaling pathway decreases resistance to bacterial pathogens. Resistance to bacterial pathogens was tested by exposing worms to *Pseudomonas aeruginosa* strain PA14 in a slow kill assay. Long-lived mitochondrial mutants, *nuo-6* and *isp-1,* have increased survival on pathogenic PA14 bacteria. Mutations affecting genes involved in the p38-mediated innate immune signaling pathway including *nsy-1, sek-1, pmk-1* and *atf-7(gof)* cause decreased resistance to bacterial pathogens in wild-type (white bars), *nuo-6* (blue bars) and *isp-1* (red bars) worms. Black dotted line indicates wild-type survival. Blue dotted line indicates *nuo-6* survival. Red dotted line indicates *isp-1* survival. Survival is measured as the number of days from exposure to PA14 (day 3 of adulthood) until death. Error bars indicate SEM of survival of individual animals. This bar graph is a summary of data shown in Figure 2 to facilitate comparison across all strains. Raw data and total N per strain are provided in **Table S2**.

**Figure S9.**
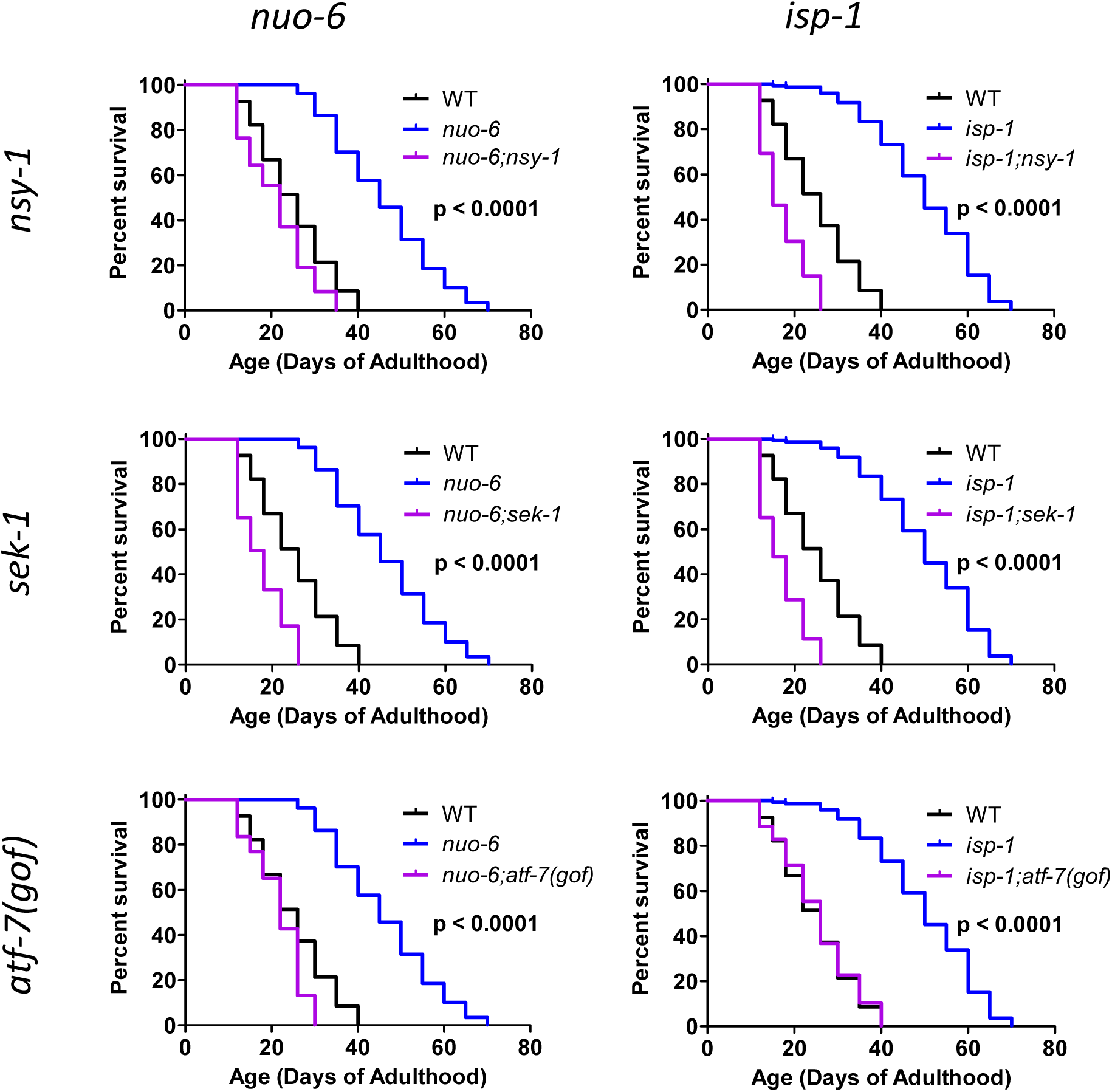
Disruption of genes involved in the p38-mediated innate immune signaling pathway abolishes the extended longevity of long-lived mitochondrial mutants independently of bacterial proliferation. Quantification of *nuo-6* and *isp-1* lifespan on non-proliferating bacteria revealed that their long lifespan is independent of bacterial proliferation. Similarly, lifespan extension in *nuo-6* and *isp-1* mutants is completely dependent on having a function p38-mediated innate immune signaling pathways as *d*eletion of *nsy-1, sek-1,* or *atf-7(gof)* completely prevented the increase in lifespan in long-lived *nuo-6* and *isp-1* mutants. Lifespans were performed in liquid culture with worms fed *ad libitum*. Bacteria proliferation was prevented through treatment with cold and antibiotics. Statistical analysis on survival plots was performed with log-rank test. p-value indicates significance of difference between blue and purple lines. All strains were tested in a single parallel experiment. Control strains are shown in multiple panels for direct comparison. Raw data and total N per strain can be found in **Table S2**.

**Figure S10.**
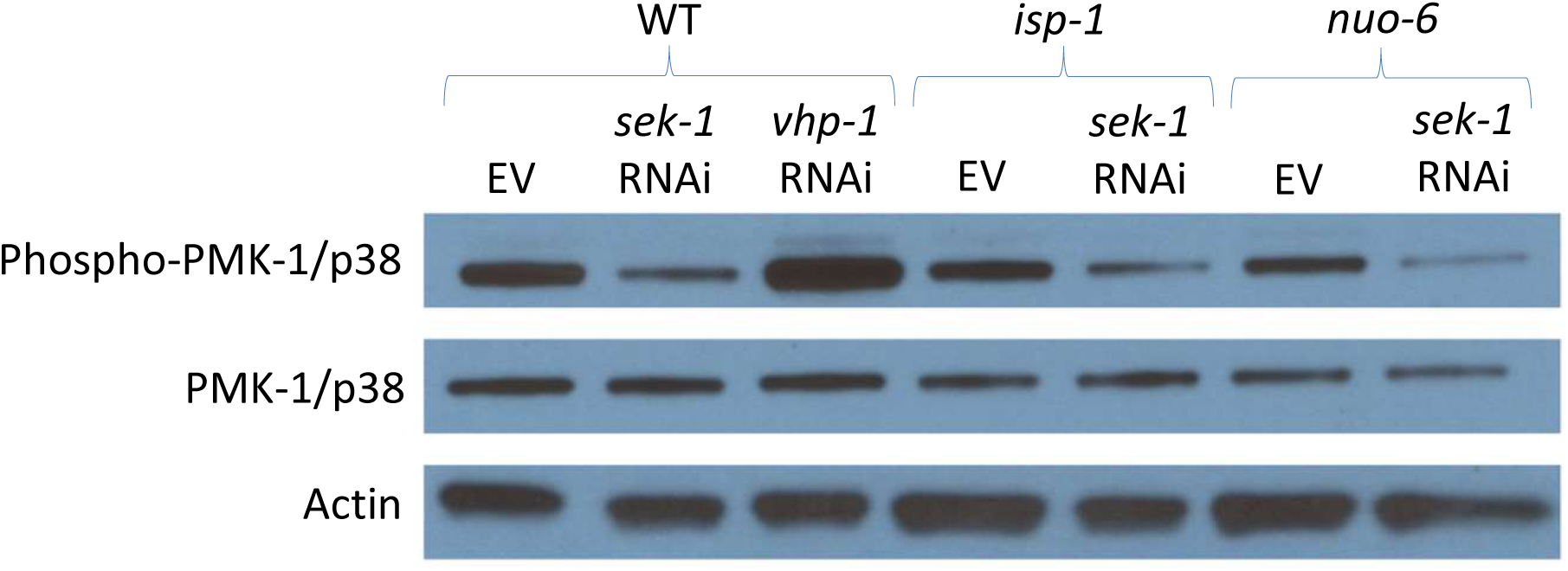
Proportion of activated PMK-1/p38 is not increased in *nuo-6* and *isp-1* worms. Activation of PMK-1/p38 was measured by levels of phosphorylated p38/PMK-1 compared to total levels of PMK-1 by western blotting. As a positive and negative controls, we examined the effect of *vhp-1* RNAi and *sek-1* RNAi. RNAi against *sek-1* resulted in decreased phosphorylation of PMK-1/p38 without affecting total PMK-1 levels. RNAi against *vhp-1* increased phospho-p38/PMK-1 levels without affecting total PMK-1 levels. RNAi treatment was begun at the L3 developmental stage and worms were collected 2 days later.

**Figure S11.**
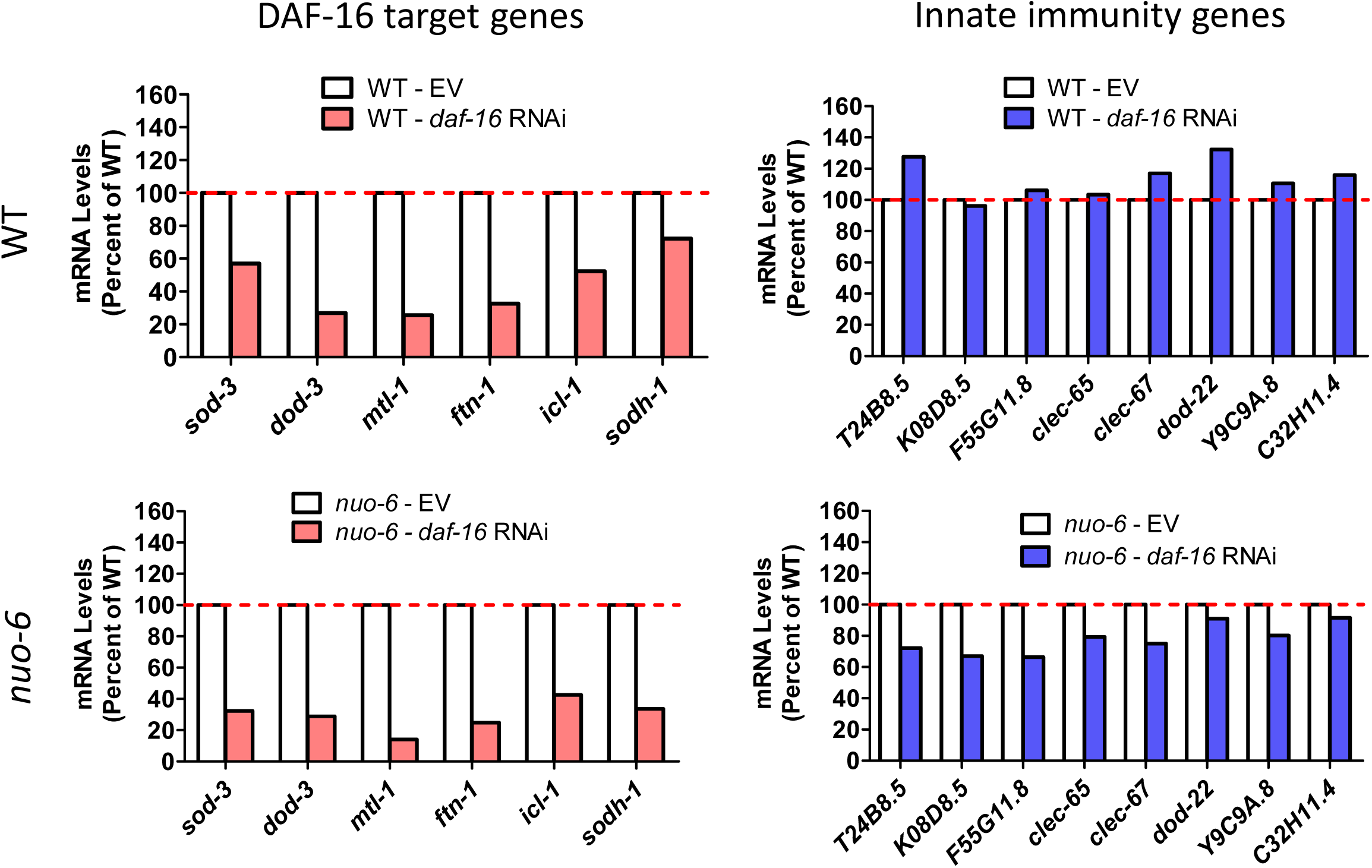
DAF-16 is not required for expression of innate immune signaling pathway target genes. *daf-16* expression was knocked down using RNAi beginning at the L4 stage of the parental generation. While *daf-16* RNAi effectively decreased the expression of DAF-16 target genes (**b**, *sod-3, dod-3, mtl-1, ftn-1, icl-1, sodh-1*) in both wild-type and *nuo-6* mutants, it did not markedly affect the expression of any of the innate immunity genes (**c**, *T24B8.5, K08D8.5, F55G11.8, clec-65, clec-67, dod-22, Y9C9A.8* and *C32H11.4*). This suggests that DAF-16 is not required for expression of innate immune signaling pathway target genes in wild-type worms and *nuo-6* mutants. RNA was isolated from six biological replicates at the young adult stage of the experimental generation. RNA from the six biological replicates was pooled for RNA sequencing.

**Figure S12.**
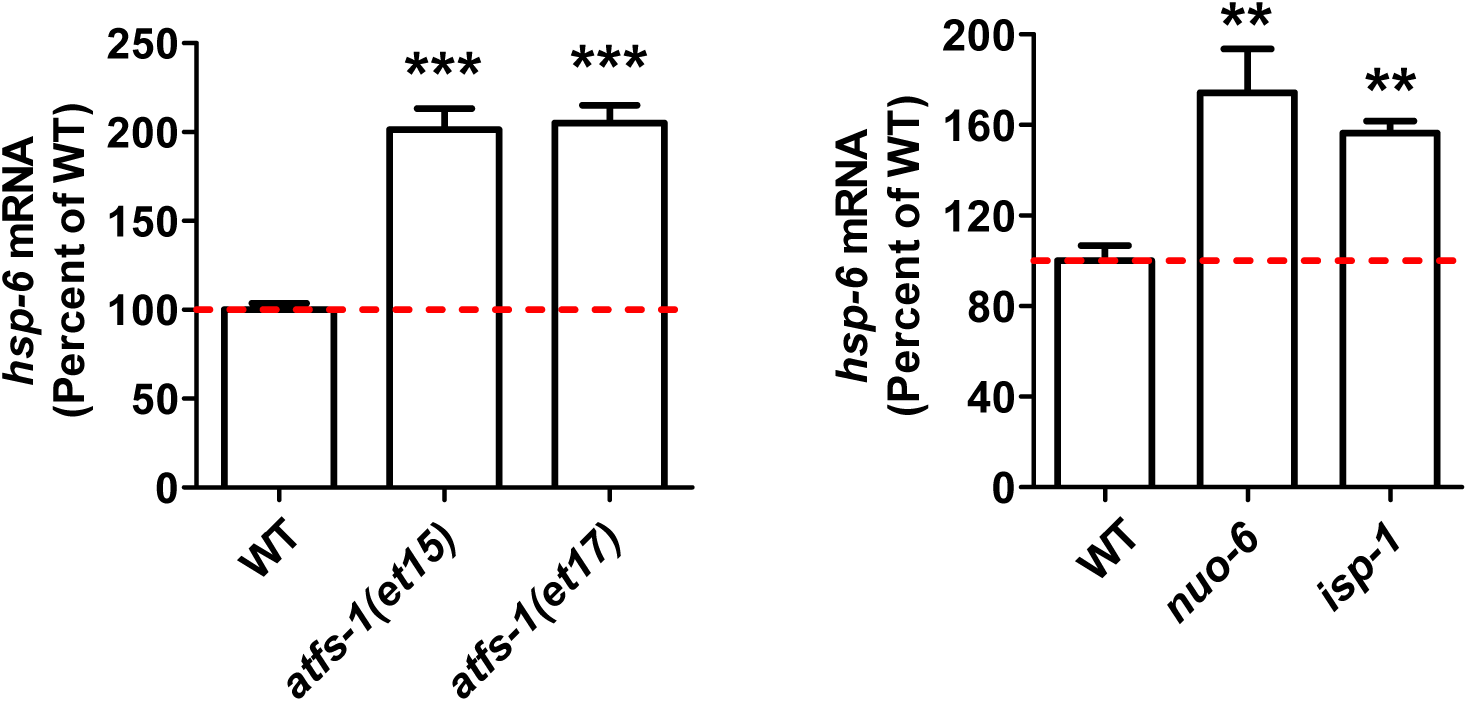
hsp-6 levels are increased in constitutively active *atfs-1* mutants. Gene expression changes were determined by RNA sequencing of six biological replicates of each genotype. Results represent counts per million (CPM) expressed as a percentage of wild-type. Error bars indicate SEM. **p<0.01, ***p<0.001.

**Figure S13.**
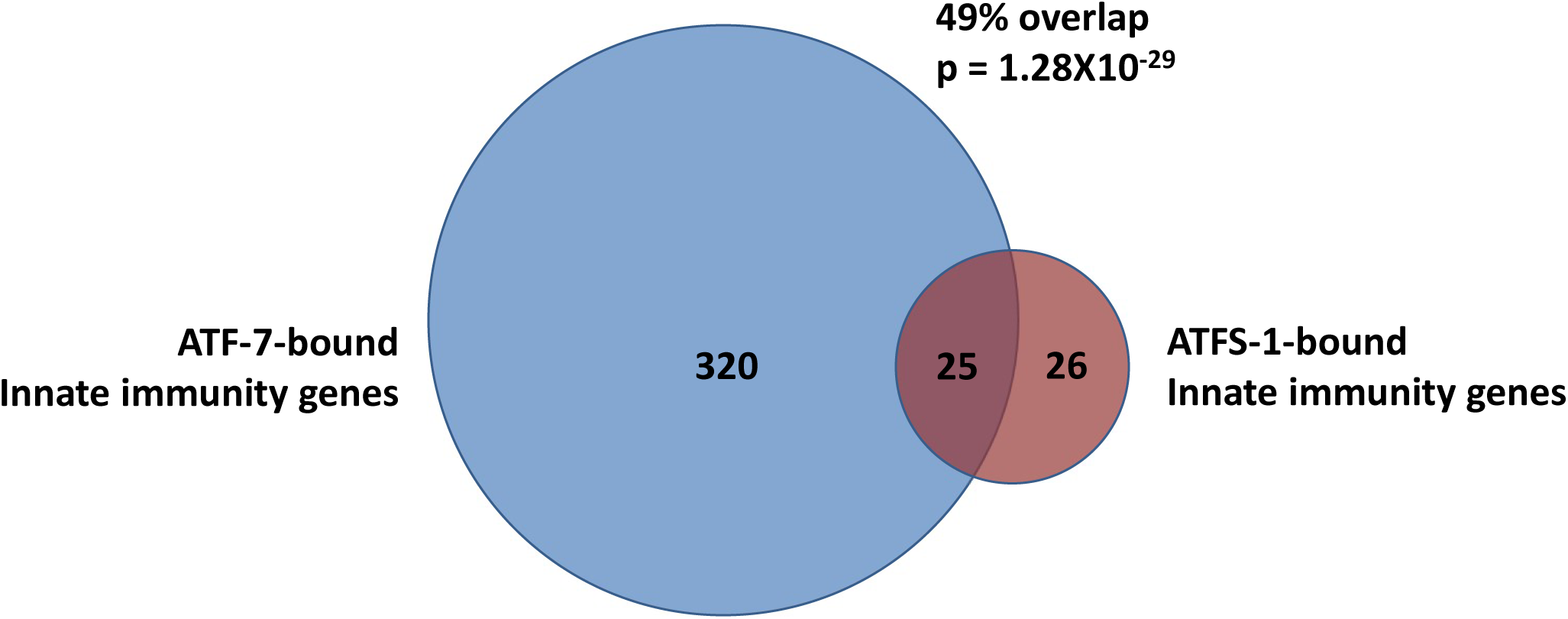
ATFS-1 and ATF-7 bind to the same genes involved in innate immunity. Innate immunity genes are defined as genes that are upregulated by a 4-hour exposure to PA14. This list was obtained from Fletcher et al., 2019. ATF-7-bound genes are genes bound by ATF-7 after exposure to PA14 from a CHIP-seq experiment conducted by Fletcher et al., 2019. ATFS-1-bound genes are genes bound by ATFS-1 after mitochondrial stress resulting from RNAi against *spg-7* as determined by a CHIP-seq experiment conducted by Nargund et al., 2015. ATF-7 was found to be bound to 345 innate immunity genes after exposure to PA14. ATFS-1 was found to be bound to 51 innate immunity genes after exposure to mitochondrial stress. Of these 51 genes, 25 were found to be in common with ATF-7-bound innate immunity genes. This highly significant overlap (p=1.28X10^-29^) clearly demonstrates that ATF-7 and ATFS-1 can bind to and regulate the same innate immunity genes. See supplemental Table 2 for the complete gene lists.

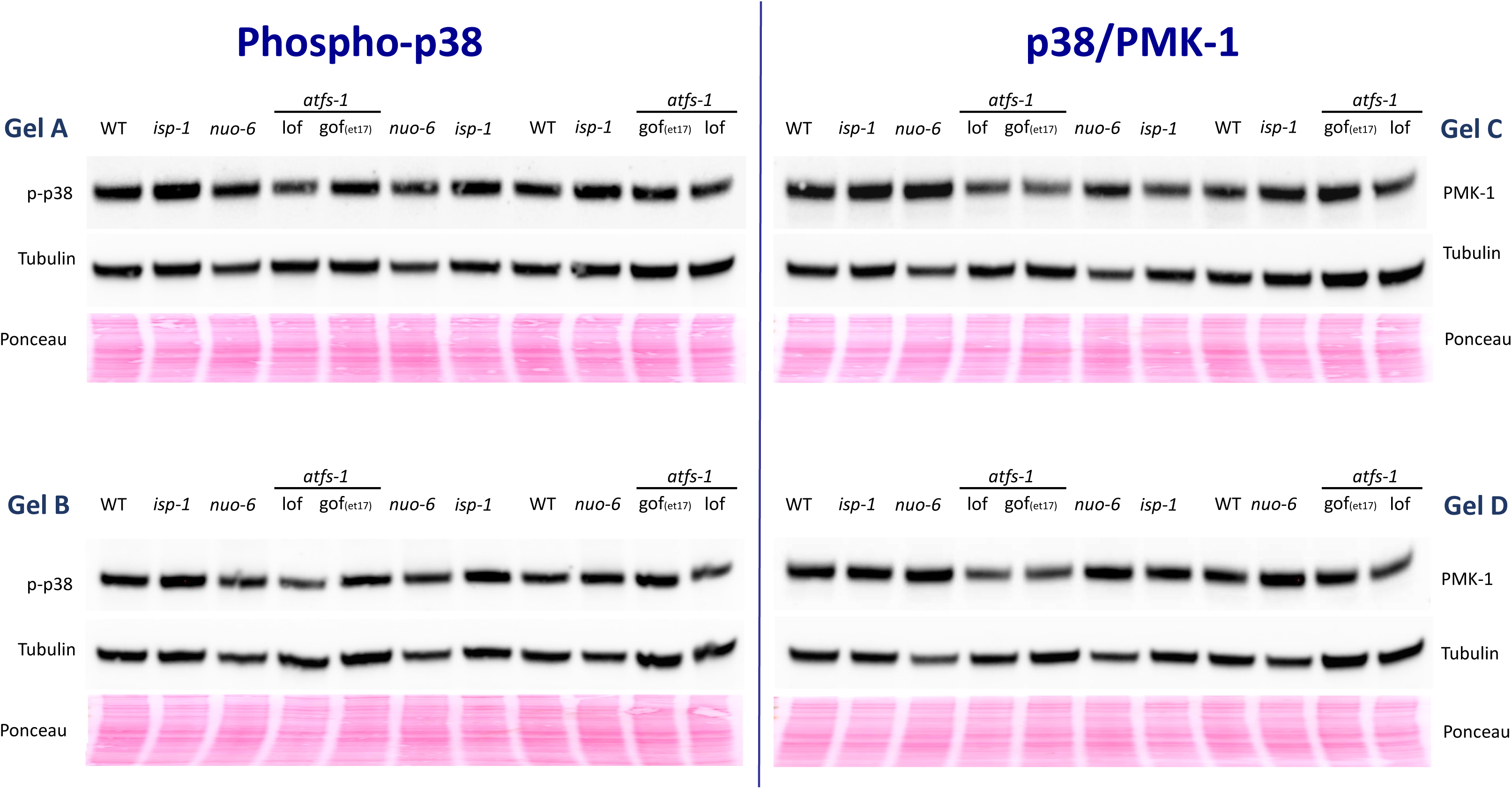

